# Engineered Protein-G variants for plug-and-play applications

**DOI:** 10.1101/2024.08.06.606809

**Authors:** Tomasz Slezak, Kelly M. O’Leary, Jinyang Li, Ahmed Rohaim, Elena K. Davydova, Anthony A. Kossiakoff

## Abstract

We have developed a portfolio of antibody-based modules that can be prefabricated as standalone units and snapped together in plug-and-play fashion to create uniquely powerful multifunctional assemblies. The basic building blocks are derived from multiple pairs of native and modified Fab scaffolds and protein G (PG) variants engineered by phage display to introduce high pair-wise specificity. The variety of possible Fab-PG pairings provides a highly orthogonal system that can be exploited to perform challenging cell biology operations in a straightforward manner. The simplest manifestation allows multiplexed antigen detection using PG variants fused to fluorescently labeled SNAP-tags. Moreover, Fabs can be readily attached to a PG-Fc dimer module which acts as the core unit to produce plug-and-play IgG-like assemblies, and the utility can be further expanded to produce bispecific analogs using the “knobs into holes” strategy. These core PG-Fc dimer modules can be made and stored in bulk to produce off-the-shelf customized IgG entities in minutes, not days or weeks by just adding a Fab with the desired antigen specificity. In another application, the bispecific modalities form the building block for fabricating potent Bispecific T-cell Engagers (BiTEs), demonstrating their efficacy in cancer cell-killing assays. Additionally, the system can be adapted to include commercial antibodies as building blocks, greatly increasing the target space. Crystal structure analysis reveals that a few strategically positioned interactions engender the specificity between the Fab-PG variant pairs, requiring minimal changes to match the scaffolds for different possible combinations. This plug-and-play platform offers a user-friendly and versatile approach to enhance the functionality of antibody-based reagents in cell biology research.

## Introduction

Antibody-based reagents play preeminently in numerous cell biology applications, including molecular detection in solutions or tissues, protein purification, ELISA assays, Western blotting, and therapeutic interventions. While these methods exhibit considerable variation in their procedures and functional readouts, they all share the fundamental reliance on antibodies as the core element. Researchers frequently depend on commercially available antibodies and design their experiments based on the accessibility of these reagents, which often can vary significantly in their quality (Couchman 2009). Additionally, a majority of these applications require the antibody to be labeled in some form. As detection reagents, antibodies may be directly labeled or necessitate the use of secondary antibodies for detection purposes (Im et al. 2019) Characterizing each antibody’s quality is imperative, yet it is often a frustrating and labor-intensive process (Schumacher and Seitz 2016). Additionally, with the transformative advancement of powerful new microscopy and proteomics platforms, the importance of high-performance antibody reagents in cell biology research has increased concomitantly.

Traditionally, immunization technologies were widely employed for antibody generation (Köhler and Milstein 1975). Although these techniques have a proven track record, the advent of recombinant versions that can be customized by molecular display technologies, including phage and yeast display methods (Boder and Wittrup 1997; Bradbury et al. 2011; Smith 1985; Viti et al. 2000), have broadened the utility of antibody-based reagents significantly. Nevertheless, significant challenges still remain to fully utilize the potential of antibody reagents because many of the most prevalent applications involve functionalizing antibodies with cargos, like tags and reporter groups, through processes that are often costly, time-consuming, and inefficient.

The most commonly employed methodologies for cargo attachment encompass chemical modifications to specific residue types or posttranslational modifications (Agarwal and Bertozzi 2015). However, these methodologies are inherently low throughput and can necessitate multiple purification steps to eliminate excess labeling agents and diminish the final product yield. These issues are further exacerbated by the fact that subsequent validation often reveals they have impaired functionality compared to their unmodified counterparts. With this backdrop, it was evident that there is a need to develop a new class of multifunctional antibody-based affinity reagents with enhanced functionality and, importantly, be highly user-friendly in their application.

With these characteristics in mind, we have developed a technology platform that utilizes a set of engineered Fab-Protein G (PG) modules that can be coupled together in multiple formats to perform complex tasks beyond what is possible with traditional antibodies. Protein G (PG) is a member of a family of immunoglobulin-binding proteins (IBP), which includes Protein A and Protein L (Björck and Kronvall 1984) and is a very robust molecule to which various types of cargo can be attached, ranging from small detector molecules to large proteins (Slezak et al. 2020). In our previous work, we used phage display mutagenesis to generate a PG variant (GA1) that had an ultra-high affinity to a set of modified Fab scaffolds with no cross-reactivity to an IgG Fc domain (Bailey et al. 2014). This provided the molecular framework whereby Fabs can bind to multiple GA1 domains that are linked together to form multivalent entities. Thus, multifunctional assemblies can be built akin to Lego blocks, pieces of which can be prefabricated as standalone units and then linked together in a plug-and-play fashion.

We have now undertaken further enhancements to the platform by expanding the number of different engineered Fab-PG pairs that are available to the researcher. A cohort of Fab scaffold types that could discriminate between specific PG variants introduces orthogonality to the system, thereby combinatorially expanding their versatility across multiple application platforms. The original Fab-PG assembly employed the PG variant, GA1, which had been engineered to recognize a non-wildtype Fab framework (Bailey et al. 2014). Thus, we felt it was important to fabricate several PG variants that recognized Fab scaffolds from different wildtype (WT) human and murine frameworks with high selectivity, enabling native human and murine Fabs to be directly plugged into the system. In principle, while the possibilities are extensive, it requires relatively few simple changes to either the Fab or PG scaffold to expand the possible unique Fab-PG pairs to facilitate highly sophisticated cell biology experiments to be performed in plug-and-play fashion. Notably, the PG scaffolds that are engineered to bind to native Fab scaffolds allow commercial antibodies also to be incorporated into the platform of applications described below.

Herein, we describe a portfolio of synthetic protein G variants exhibiting selective binding characteristics that facilitate cargo attachment to multiple Fab scaffold types. In the simplest manifestation, the orthogonality of the system allows for a group of PG variants to be used, fused through a linker to different fluorescently labeled SNAP-tags. These can then be mixed with Fabs of a chosen scaffold type in a multiplexed fashion to detect several antigens in a single experiment. We show that Fabs can be readily attached with high affinity to a module containing a PG linked to a dimer Fc domain to produce a plug-and-play IgG-like assembly. The utility of the system can be further expanded to produce bispecific analogs using the “knobs into holes” strategy by matching two different pairs of Fab-PG Fc variants to produce plug-and-play bispecific antibodies in one step. We describe how these bispecific modalities form the building block for fabricating potent Bispecific T-cell Engagers (BiTEs) and show their potency in cancer cell-killing assays. We also delineate how the PG variants that were originally engineered to be Fab-specific were back-engineered to reintroduce Fc binding mutations, converting them into Fc specific binders. This allows for the coupling of cargos in a similar fashion to the Fc domain of IgGs, which can then be manipulated as components in the plug-and-play assemblies in a similar fashion to their Fab scaffold counterparts.

While the platform described offers many possibilities by combining the various component parts, we show through crystal structure analysis that a surprisingly few strategically positioned interactions engender the specificity between the Fab-PG variant pairs. Thus, matching the scaffolds for the different possible combinations requires very few mutations and virtually no high-level molecular biology expertise. In fact, the vast majority of the applications can be performed using only a couple of Fab scaffolds and two PG variants. Importantly, the PG constructs are easy to produce and can be stockpiled frozen or lyophilized to be used as off-the-shelf reagents. Thus, this plug-and-play protein G-base antibody platform will provide researchers with an ease-of-use tool kit that greatly simplifies many challenging cell biology applications and increases the throughput significantly.

## Results

### Development of new Protein-G variants with Fab scaffold-based selectivity

In previous work, an ultra-high affinity Fab scaffold (Fab^LRT^) was developed that binds (100 pM) the GA1 variant of Protein-G (PG) (Slezak et al. 2020). However, GA1 was originally generated for a mutated version of the Herceptin Fab scaffold (Fab^S^), it did not bind to wild-type versions of the scaffold or any murine orthologs. More universal versions of GA1 could have significant value by allowing the plug-and-play capability of the system to be extended to facilitate applications exploiting multiplexing between other Fab scaffold types. Two categories of PG variants were envisioned. One would be fully universal being able to bind all native scaffolds broadly. The second category would bind with high affinity to a wild-type (*wt*) human Fab (Fab^H^), but not to Fab^LRT^. This requirement would provide an orthogonal pair that could be used simultaneously in experiments that required the delivery of two different cargos.

## 1) Generation of customized Protein-G binders to different Fab scaffolds

To generate these scaffold-specific Fabs, we employed phage display mutagenesis using the GA1 variant as the starting point. Phage display libraries were designed focusing on the C-terminal cap of the α-helix in the heavy chain (Hc), central in the interaction with the constant portion of the Fab light chain (Lc) (Figure 1A, B). Figure 1C provides the sequences of the different Fab scaffold types at their point of engagement to GA1 (residues 123-127). Using Kunkel mutagenesis, we generated six libraries based on a hard randomized strategy (NNK diversity) covering residues 38-43 in the α-helix region of GA1 (Figure 1D). Four of the libraries preserved His at position 42, since it had previously been found to improve the pH dependence of the engineered GA1 (Bailey et al. 2014). To enable possible conformational change in the helix, randomization of several residues bordering the helix that contacted the Fab Lc was introduced. The phage display selections involved using biotinylated Fabs as the target antigens.

**Figure 1.**
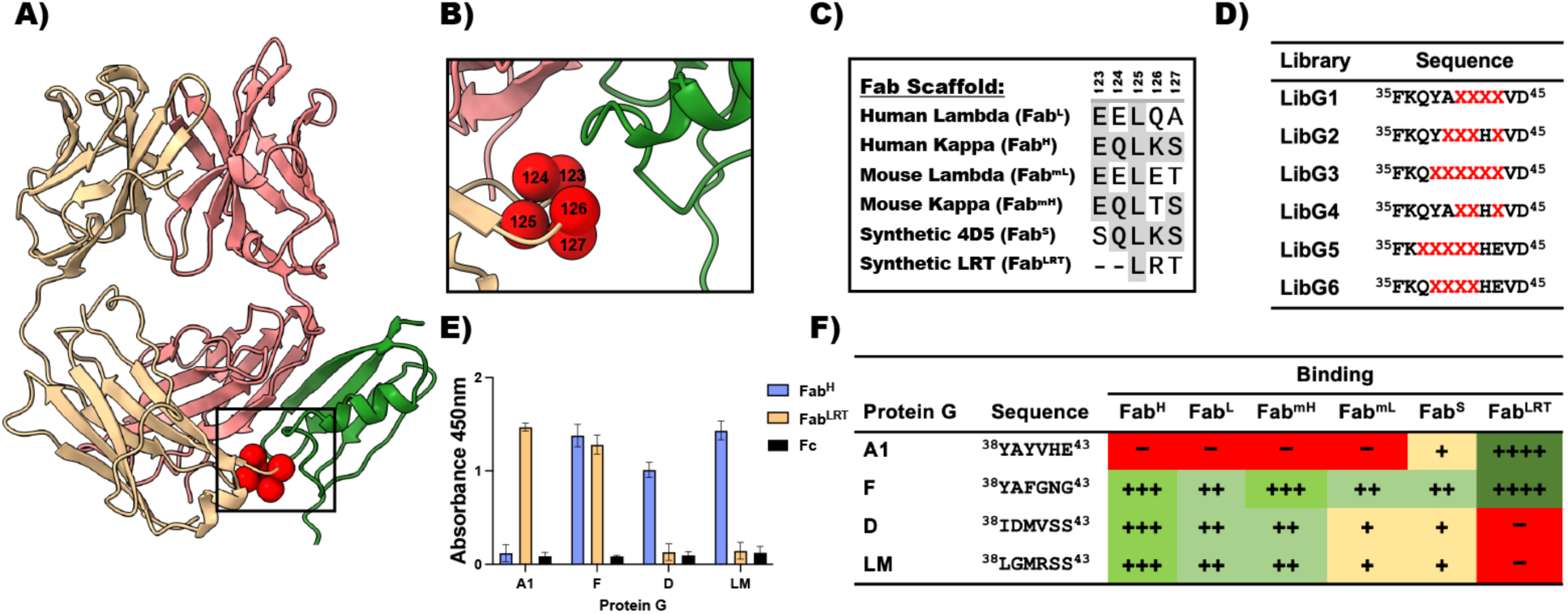
New Protein G engineering. (**A**) The Interface between constant part of Fab Light Chain and Protein G. Main interaction comes from an antiparallel *β*-strand configuration of Fab Hc and protein G. Additionally, protein G interacts with Fab Lc via a *α*-helical cap. Spares represent randomized residues in the constant part of Fab Light Chain. Fab Heavy Chain is colored in red, while Light chain is colored as yellow. (**B**) The interaction between Fab Light Chain and Protein G is limited to 5 amino acids. Naturally existing Fab scaffolds contain conserved glutamic acid at position 123 and leucine at position 125. Interestingly, GA1 was previously engineered against Fab^S^, and does not recognize any of the naturally existing Fab scaffolds due to a negative charge at position 123. (**C**) Sequence alignment of constant part of the light chain (position 123-127) in different Fab scaffolds. The sequence alignment of Fab area recognized by protein G shows the opportunity for affinity improvement. (**D**) Generated phage libraries where protein G helical cap interacting with Fab light chain is randomized. Hard randomization (NNK) of selected residues is represented by “X”. (**E**) ELISA of selected protein G variants against Fab^H^, Fab^LRT^ and Fc. Results show the high specificity towards certain Fab scaffolds that allows formation of orthogonal pairs that would not cross-react. For example, GA1-Fab^LRT^ and GD-Fab^H^. Protein GF is a universal high-affinity Fab binders. Importantly, the interaction with the FC portion of the IgG has been engineered out. (**F**) The selected sequences of engineered protein G variants. The binding affinities to different Fab scaffolds are indicated with “+”. No binding is indicated with “-“. Sequences of all the generated protein G variants are listed in the supplementary materials.

### 1.1). Generation of the universal Protein-G variant

Five rounds of phage display selection were performed to generate a universal PG variant, sequentially swapping different Fab scaffolds from round to round. The 1^st^ round targeted Fab^H^ (*wt* Herceptin scaffold having a kappa (κ) light chain), followed by Fab^L^ (human lambda (λ) Lc), Fab^LRT^, and Fab^S^ in successive rounds, ending with the final round repeating with the Fab^H^ antigen. Additionally, the antigen concentration was systematically reduced to increase stringency, starting with 1 µM in round 1 and ending with 1 nM in the final round. A phage ELISA was performed on 96 clones to validate the specificity of the selected PG variants. This resulted in the identification of seven universal PG variant candidates. Interestingly, a majority of the universal PG variants retained the *wt* PG composition at positions 42 and 43 (Figure S1). This selection resulted in the universal protein GF variant (Figure 1 E, F).

### 1.2). Generation of Fab^H^ PG variants

Next, we generated a set of PG variants recognizing wild-type κ and λ Fab scaffolds (Fab^H^), but not the engineered ones (Fab^S^, Fab^LRT^). Fab^H^ is a wild-type human (Herceptin) Fab with a κ light chain. The selection aimed to produce a PG variant that bound to human *wt* Fab scaffolds with both κ and λ Lc, but not the modified Fab^LRT^ scaffold. Five rounds of selection were performed using Fab^H^ as the target antigen. To counter-select against binders to the Fab^LRT^ scaffold, 2 mM of unbiotinylated Fab^LRT^ was added to the selection buffer in each step of the biopanning. Thus, potential Fab^LRT^ binders were captured and eliminated in the wash steps, effectively depleting them from the library. The antigen concentration was systematically reduced round to round, starting with 1 mM and ending with 1 nM. After the final round of selection, phage ELISA was performed. The positive clones were sequenced, resulting in five unique high affinity Fab^H^ binders with no measurable cross reactivity to Fab^LRT^. Sequence alignment of these variants shows diversity was introduced in each position of PG with a dominating methionine at position 40 (Figure S1). Notably, although selections were made with a Fab having a κ light chain, both the GD and GLM variants showed varying cross-reactivity towards human and murine Fabs with λ light chains (Figure 1 E).

### 1.3). Biophysical characterization of the variant PGs

Surface plasmon resonance (SPR) was performed to determine the binding affinity and kinetics of the variant PGs. SPR showed a significant improvement in affinity and specificity compared to *wt* PG (Figures S2, S3). The generation of multiple universal PGs was confirmed by testing the binding with the most common types of Fab scaffolds: Human κ (Fab^H^), Human λ (Fab^L^), Murine κ (Fab^mH^), and Murine λ (Fab^mL^), respectively) (Table 1). The most promising universal PG candidate, Protein-GF, exhibits high affinity towards all tested Fab scaffolds – Fab^H^ (K_D_-1.9 nM), Fab^L^ (K_D_-21 nM), Fab^mH^ (K_D_-3.2 nM), Fab^mL^ (K_D_-37 nM), and Fab^LRT^ (K_D_-0.9 nM). Additionally, the results indicate the successful generation of the orthogonal set of PG-Fabs that would allow for simultaneous usage with the Fab^LRT^-GA1 platform. Several generated PG variants are characterized by a high affinity and specificity towards Fab^H^ (Figure 1E) with the best candidates (GD, GLM) having affinities of 6.4 and 8.9 nM, respectively (Table 1). The high specificity of the system was evaluated by ELISA, injecting 100 nM of Fab^LRT^ over the GD and GLM, confirming no detectable binding in either case (Figure 1E). Both GD and GLM recognized Fab^L^. However, the affinity was ~3-fold lower than Fab^H^, which is consistent with the decreased affinity observed for GF and Fab^L^ (Table 1). A compilation of the specificity of PG variants for each scaffold type is presented in Figure 1F.

**Table 1.** Binding affinities of engineered protein G to different Fab scaffolds measured by SPR. All SPR sensograms are shown in supplementary material.

| Protein G | Fab | $K_{on} (M^{-1} s^{-1})$ | $K_{off} (s^{-1})$ | $K_D (nM)$ | $\chi^2 (RU^2)$ |
| --- | --- | --- | --- | --- | --- |
| **D** | **Fab<sup>H</sup>** | $8.3 \times 10^5$ | $5.3 \times 10^{-3}$ | 6.4 | 1.3 |
| | **Fab<sup>L</sup>** | $1.3 \times 10^5$ | $2.1 \times 10^{-3}$ | 16 | 0.7 |
| **LM** | **Fab<sup>H</sup>** | $2.6 \times 10^5$ | $2.2 \times 10^{-3}$ | 8.9 | 0.7 |
| | **Fab<sup>L</sup>** | $1.4 \times 10^5$ | $3.0 \times 10^{-3}$ | 21 | 2.1 |
| **F** | **Fab<sup>H</sup>** | $4.3 \times 10^5$ | $8.2 \times 10^{-4}$ | 1.9 | 0.2 |
| | **Fab<sup>mH</sup>** | $4.4 \times 10^5$ | $1.4 \times 10^{-3}$ | 3.2 | 0.4 |
| | **Fab<sup>LRT</sup>** | $5.2 \times 10^5$ | $4.9 \times 10^{-4}$ | 0.9 | 0.3 |
| | **Fab<sup>L</sup>** | $1.5 \times 10^5$ | $3.2 \times 10^{-3}$ | 21 | 1.0 |
| | **Fab<sup>mL</sup>** | $1.3 \times 10^5$ | $4.8 \times 10^{-3}$ | 37 | 1.7 |

## 2) Structural insight into specificity differences between the protein GF and GD

The crystal structures of an antigen protein complexed with GF-Fab^H^ and GD-Fab^H^ described below were undertaken to establish the structural elements conferring the PG variants’ individual specificities. The GF variant is “universal” in binding to all Fab formats. This contrasts with the GD variant, which does not bind to the engineered Fab^LRT^ scaffold, but all the natural human and murine scaffolds. In both cases, the target antigen protein used in the structure determination was the histone chaperone Anti-silencing factor 1 (ASF1) protein (Bailey, Sheehy et al. 2018). The full complex contained the common components ASF1 and anti-ASF1 Fab^H^ combined with either GF or GD to form a tri-component complex. The anti-ASF1 Fab used was Fab E12 from the previous structure study (Bailey et al. 2018), but with its framework modified from Fab^S^ to Fab^H^. It is designated Fab E12^H^ for this set of structure determinations. The experimental particulars are provided in Table S1.

### 2.1). ASF1:Fab E12^H^:GF

Protein G interacts with the Fab scaffold through an antiparallel β-strand interaction with the heavy chain and a set of helical cap interactions with the light chain (Figure 2A). Since our protein engineering strategy did not involve altering this β-strand contact, the H-bonding in the strand was identical to the previously analyzed GA1-Fab^LRT^ structure (Slezak et al. 2020). The main area of difference was the PG’s helical cap interaction with Fab Lc, since the helical cap was broadly randomized in the PG phage display libraries. The Fab E12^H^-GF interface is formed through two contacts that bury ~530Å^2^ through its interaction with Hc and ~198Å^2^ with Lc. Based on the buried surface area, most of the GF interaction surface emanates from the contact with the Fab’s heavy chain, which includes an H-bond formed between Y38 from GF and P124 of the Fab (Figure 2B). A further set of interactions is formed between Y38, G41, and N42 from GF with S125, F127, and V212 from the Fab Hc (Figure 2B). Notably, these heavy chain residues have conserved sequences among all the Fab frameworks, explaining the universality of the GF interactions. The structure analysis also revealed a close interaction with the side chain of F40 to the Fab Lc, as described previously (Slezak, Bailey et al. 2020).

**Figure 2.**
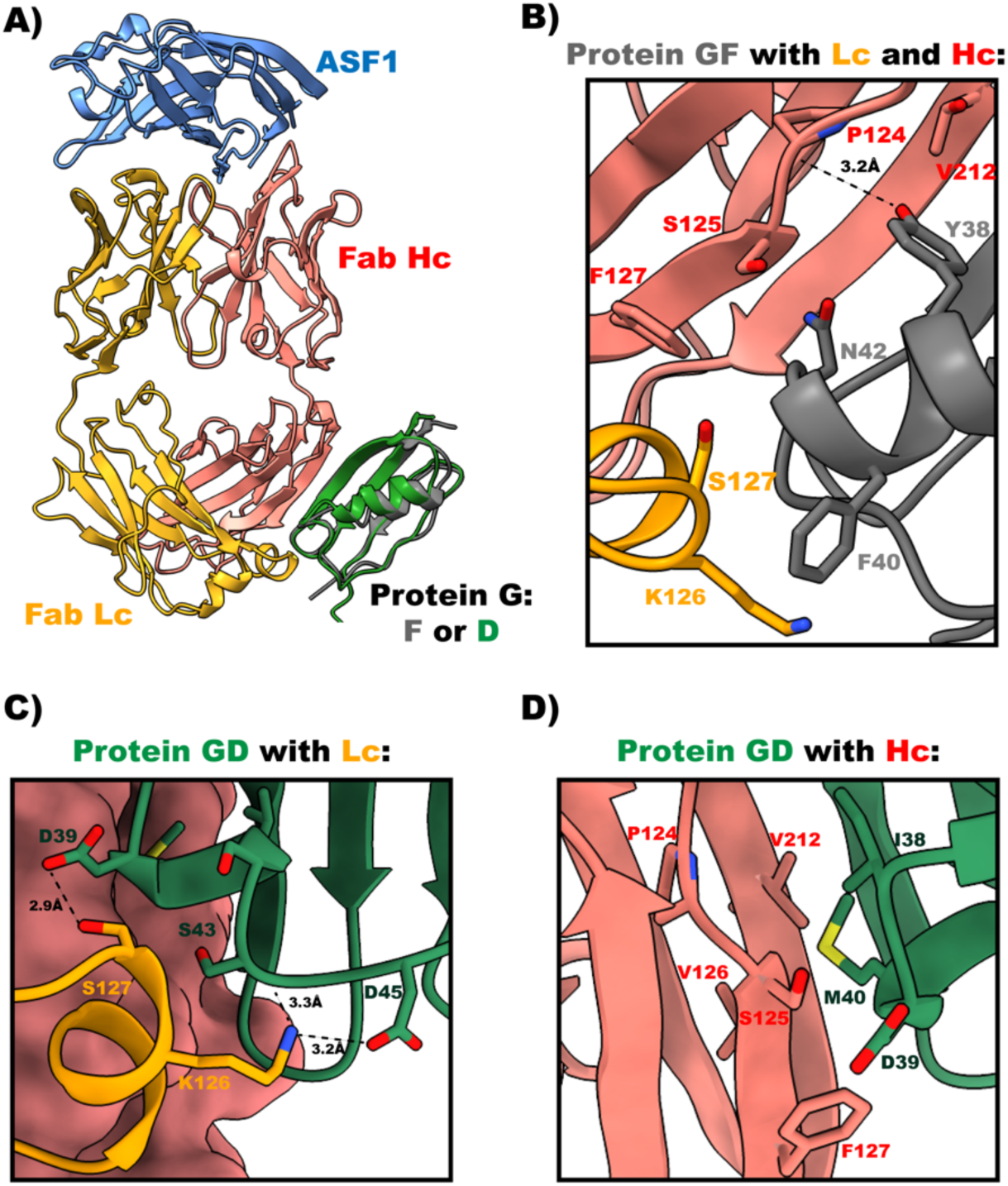
The structure of the ASF1-E12 Fab^H^ with Protein GF and Protein GD. (**A**) General view of the structure. (**B**) Interface between Protein GF and Fab^H^. Universal binding of GF comes from the extensive hydrophobic interactions of Y38, G41 and N42 with the Fab Hc. Additionally, Y38 forms a hydrogen bond with a main carbonyl of P124 from the Fab Hc. F40 placed itself between K126 and S127, forming the only interaction of GF with the Fab Lc. (**C**) GD interaction with Fab Lc. The specificity of GD is driven by the set of H-bonds formed by D39, S43 and D45, that is disturbed by the significant loop rearrangement caused by the two amino acid deletion in Fab^LRT^. Side chains of S127 from Fab^H^ form hydrogen bonds with D39 from GD. Additionally, K126 forms two hydrogen bonds with a side chain of D45 and a main chain carbonyl of S43 from GD. (**D**) GD interaction with Fab Hc. I38, D39 and M40 are engaged in several hydrophobic interactions with Fab Hc. M40 is placed in the hydrophobic pocket formed by P124, S125, V126 and V212 from the Fab Hc. I38 and D39 interact with S125 and F127 from Fab Hc, respectively. Molecules are colored as follows: ASF1-blue, Fab Hc-red, Fab Lc-orange, Protein GF-gray, Protein GD-green. PDB: (9AVO, 9AWE).

### 2.2). ASF1:Fab E12^H^:GD

The GD variant is more selective than GF because it does not bind Fab^LRT^. The Fab^H^-GD interface is formed through two principal contacts that bury ~596Å^2^ for interaction with Hc and ~185Å^2^ with Lc (Figure 2C, D). The specificity of GD comes from the set of hydrogen bonds formed by D39, S43, and D45 from the GD with Fab^H^ Lc. Side chains of S127 from Fab^H^ form H-bonds with D39 from GD. Additionally, K126 forms two H-bonds with a side chain of D45 and a main chain carbonyl of S43 from GD (Figure 2C). These interactions are eliminated by the significant loop rearrangement caused by the two amino acid deletions in Fab^LRT^ (Figure S4B). Interactions of I38, D39, and M40 from GD are formed with Fab Hc. M40 of GD is buried at the Hc Fab interface formed by P124, S125, V126, and V212 (Figure 2D). Additional interactions between I38 and D39 of GD with S125 and F127 from Fab Hc are formed, respectively.

Taken together, this structural information provides insight for the basis of the PG variant specificities. In the case of variant GA1, the inability to bind to Fab^H^ or other wt-Fab Hc frameworks can readily be attributed to the charge repulsion between E43 of GA1 and E123, which is present in all wt-Fab Hc molecules (Figure S4A). The inability of Fab^LRT^ to bind to the GD variant can be directly attributed to the two amino acid deletion in the Fab^LRT^’s light chain, which induces a significant loop rearrangement which places its T127 side chain into steric conflict with D39 of the GD variant (Figure S4B).

## 3) Applications exploiting the modular design of PG variants

The Protein G-Fab plug-and-play system described below was developed to provide researchers with a highly diverse toolkit of antibody-based reagents that can be readily applied across many different types of applications. This system overcomes many of the limitations that have plagued the general use of traditional antibodies in some complex applications. The basis of the plug-and-play system is that highly customized affinity reagents can be constructed from simple component parts that can be assembled like Lego blocks by taking advantage of the ultra-high affinity protein Gs with their orthogonal Fab scaffolds. Thus, multifunctional assemblies that would be highly challenging to produce as single entities can be readily fabricated with interchangeable parts that easily alter specificity and avidity to match the particular application.

### 3.1). Stability of Plug and Play PG complexes in cell-based applications

The modular design that allows for the ability to “snap-on” functional cargo onto antibodies and antibody fragments in a facile manner has obvious applications and is advantageous over traditional chemical coupling methods and protein fusions. This modular design concept also provides easy multiplexing, allowing for high throughput processing of many samples with minimal system changes. One consideration, however, is that the attachment is non-covalent and there might be a concern that the cargo could disengage during the experiment’s lifetime. To better quantify the degree of dissociation under typical experimental conditions, we undertook a time course analysis of the stability of a typical PG variant-Fab complex. This involved measuring the changes in binding on HeLa cells of an EGFR Fab with a human kappa scaffold (Fab^H^) premixed in a 1:1 molar ratio with the PG variant, GLM, that was labeled with Alexa647. The cells were rigorously washed and the binding of the EGFR Fab was visualized using Flow Cytometry over a time interval of 24 hrs. The cells were kept at 4°C during the experiment to minimize receptor internalization. Figure S5 shows the average MFI observed over six time points from 0 mins to 24 hrs is virtually unchanged over the full-time course. For this analysis, we selected a Fab-PG variant combination (Fab^H^-GLM) that was one of the lower affinity pairs (9 nM). Most of our cell biology applications utilize the Fab^LRT^-GA1 pairing, which has an affinity about 100-fold higher (~ .1 nM). Thus, even using the lower affinity pair, the plug-and-play module is as functionally effective as the covalent alternatives during the time course of most typical cell biology experiments.

### 3.2). Generation of the Fc specific Protein-G variant

Many cell based experiments utilize commercially available antibodies in IgG format to exploit avidity effects, cell killing properties and detection by off-the-shelf secondary antibodies. Thus, we endeavored to develop a PG scaffold that could be used in a plug-and-play fashion with any full-length antibody. This allows myriad cargo types to be facilely attached to any commercial human IgG. *Wt* Protein G binds to IgG molecules through both the antibody’s Fab and Fc domains. For most applications, it is important to separate the binding between the Fab and Fc domains to eliminate undesired cross-reactivity that might interfere with interpretations in many cell based experiments. The PG variants described above were engineered to eliminate any binding to an IgG Fc domain. To accomplish this, the original GA1 variant from which the specificity matured GF, GD and GLM variants were derived, included a set of mutations introduced to specifically inhibit binding to the Fc portion of an IgG antibody. To convert GA1 into an Fc specific binder, a reverse engineering strategy was used. Based on the previously published structure of the *wt* protein G in complex with the Fc domain of human IgG (Sauer-Eriksson, Kleywegt et al. 1995), we selected seven residues in GA1 that had originally been mutated to eliminate Fc binding and reverted their sequences back to *wt* PG (Figure 3A). Three separate variants were created by making further mutations in the helix cap region of PG. Binding kinetics were determined by SPR, showing the successful generation of G-Fc, which possesses a low nM affinity to the Fc domain (3.8 nM) with a prolonged dissociation rate (Figure 3B). A 200 nM injection of Fab^H^ tested the high specificity of the G-Fc by showing no detectable binding to the Fab was observed (Figure 3C). Variants G-Fc2 and G-Fc3 displayed somewhat lower binding affinities and less optimal kinetics; thus, all our experiments were performed with the G-Fc variant.

**Figure 3.**
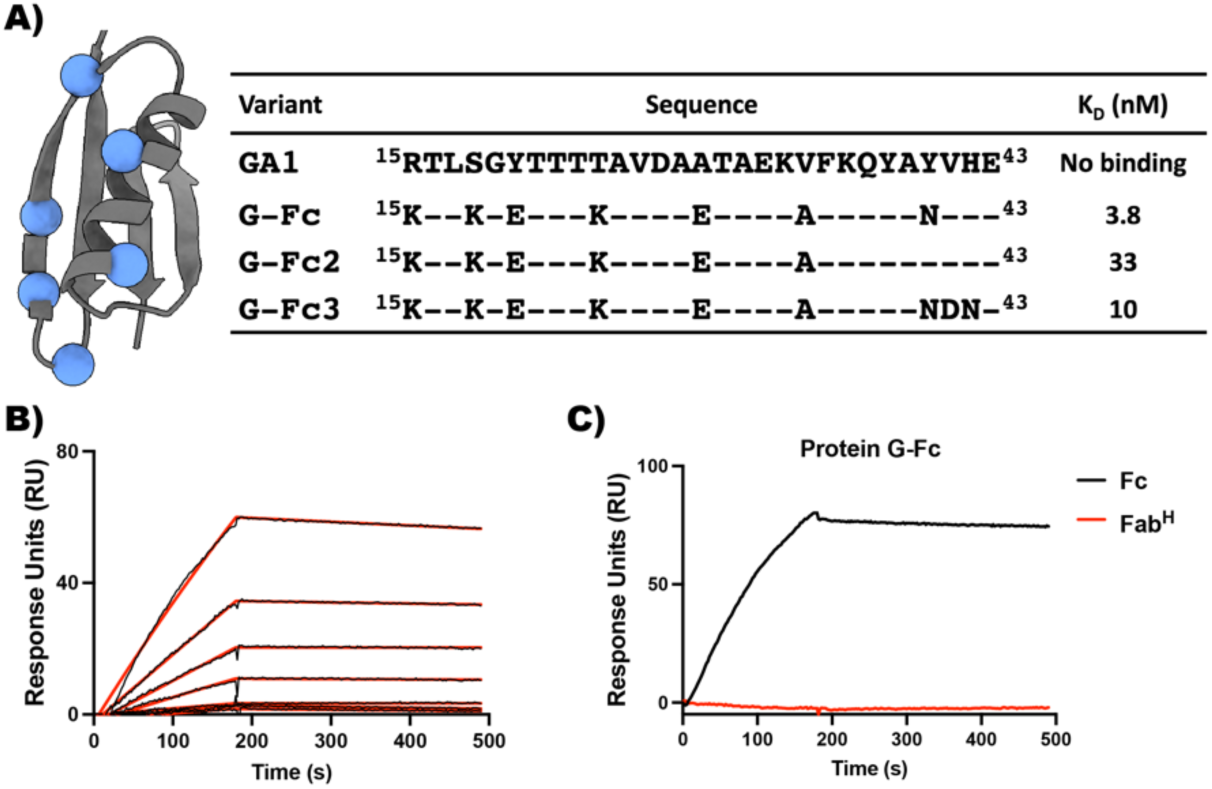
Engineering of Protein G-Fc. (**A**) The sequences of Protein G-Fc variants. Residues mutated to generate a Protein G-Fc are represented as blue spheres. (**B**) SPR sensogram showing the interaction of G-Fc with human Fc. For the kinetic experiment, analytes were serially diluted two-fold, starting at 100 nM. (**C**) A single injection of Fc and Fab^H^ on G-Fc. 200 nM of Fc and 200 nM of Fab^H^ were injected, and no binding to Fab^H^ was observed.

### 3.3). Plug-and-play IgG

Due to their avidity effect, enhanced binding affinity is one advantage of bivalent IgGs over monovalent Fabs. In this regard, fusing together multiple GA1 domains like beads on a string can be exploited to increase the binding of cargo to immobilized targets, as shown in Figure S6, where both the affinity and kinetics are substantially improved going from mono- to bi- to tri-valent modalities: 15-fold, 50-fold, respectively. The constructs are very versatile in that the number of domains and linker lengths between them are easily modified and thus, can be customized to the particular application. While the ability to use linked GA1 domains systematically increases the affinity of the construct to its designated target, it lacks the functional attributes innate in Fc domains of natural antibodies. Thus, we thought that a significant enhancement would be to fabricate a plug-and-play IgG-like molecule. Conceptually, this could facilitate the facile interchange of Fab components through the PG linking technology while having the advantages of the structure and function of natural antibody molecules.

The basic PG-IgG framework fabrication is straightforward and retains all elements of a natural antibody. The construction involves fusing two PG arms to an IgG Fc base; the PG arms can be any of the variants described above to provide a panel of different specificities. Fabs having a particular PG specificity can be simply introduced to mimic the full-length bivalent IgG (Figure 4A). The linker lengths between the PG domains and the Fc fragment can be adjusted to optimize the molecule’s efficacy with respect to a particular application. The initial test of this assembly was performed using two GA1s fused to a human Fc domain via a 17 residue linker. This linker length is similar to that found in natural antibodies. A conformation-specific anti-MBP Fab^LRT^ (7O Fab^LRT^) was associated with the Fc through its GA1 arm to detect extracellular MBP that had been stably expressed on the surface of the HEK cell line (Figure S7). An attribute of using the conformational specific 7O Fab was that it also provided an adjustable binding switch that could be controlled by systematically altering the concentration of maltose. Using an ELISA assay, we determined that 7O in its Fab format in the absence of maltose bound to cells with an EC_50_ value of 10 nM compared to ~3 nM in the GA1-Fc bi-valent format, demonstrating the bivalent avidity effect. In the presence of 10 μM maltose, no binding was observed for the monovalent Fab compared to 85 nM for the GA1-Fc format (Figure S8)

**Figure 4.**
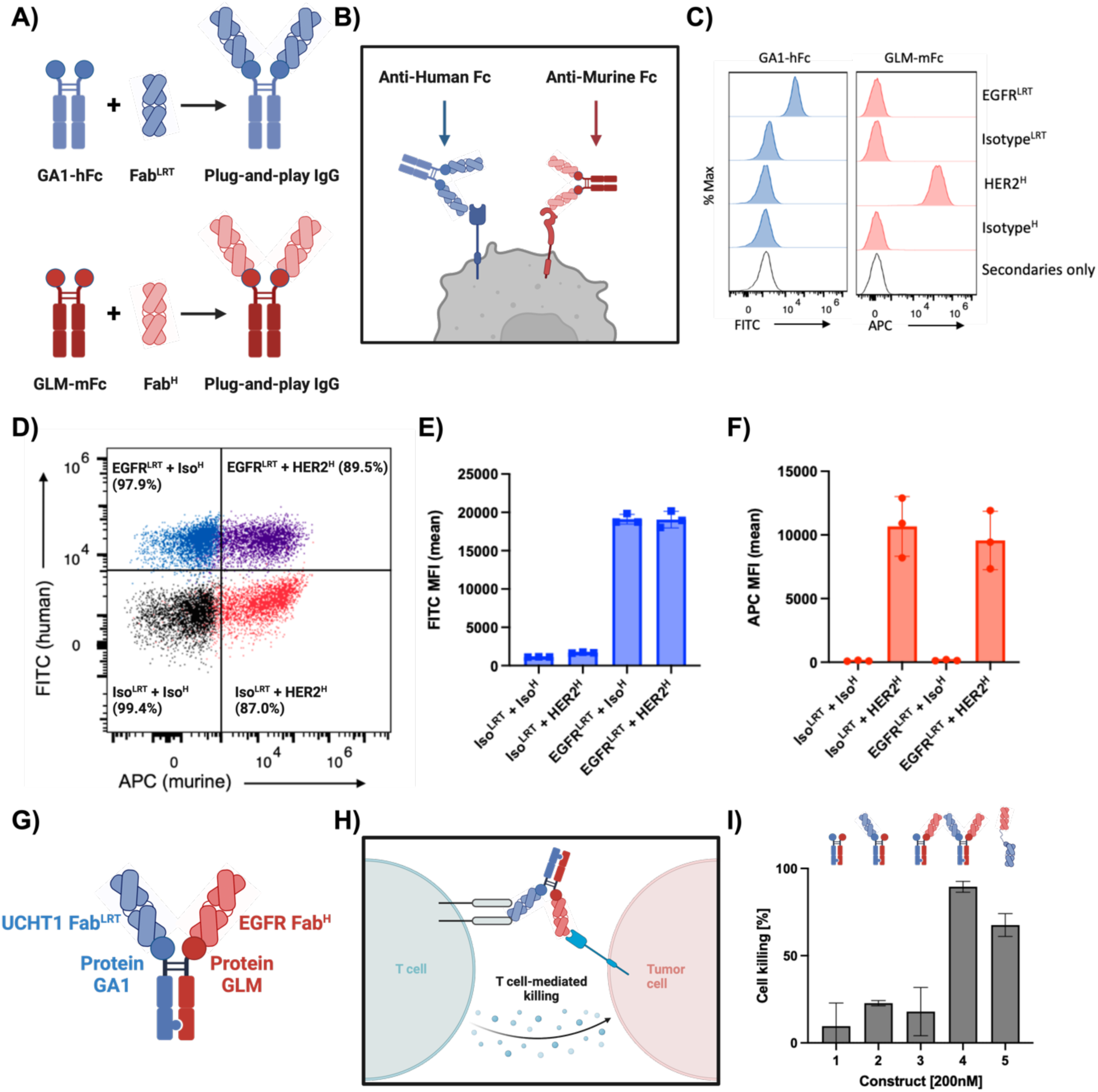
Protein G-Fc fusion enables modular assembly of bivalent IgG-like sABs. (**A**) Schematic of plug-and-play IgG assembly with human or murine Fc using orthogonal protein G-Fab scaffold pairs (**B**) Orthogonal pG-Fc fusions enable simultaneous detection of two different sABs binding to the cell surface. Model for secondary detection by anti-Fc secondary antibodies that recognize either human IgG1 Fc or murine IgG2a Fc. (**C**) HCC1954 cells were stained with IgG-like sABs targeting EGFR (Fab^LRT^ format) or HER2 (Fab^H^ format). *Left panel*, anti-human-Fc-A488 (FITC) recognizes only the combination of GA1-hFc + EGFR^LRT^. *Right panel*, anti-murine-Fc-A647 (APC) recognizes only the combination of GLM-mFc + HER2^H^. (**D**) Simultaneous staining of EGFR and HER2 using the IgG-like assemblies shown in (C). (**E**) Quantification of mean fluorescence index (MFI) in the FITC channel to visualize EGFR^LRT^ binding. (**F**) Quantification of MFI in the APC channel to visualize HER2^H^ binding. (**G**) Bispecific Easy-to-Use plug and play IgG-like Molecule (BEZ-IgG). Schematic of plug-and-play Bispecific IgG-like assembly. “Knobs-into-holes” heterodimerization technology allows for the generation of Fc-fusion dual protein G molecule which contains engineered protein GA1 and GLM that allows for the specific recognition of Fab^LRT^ or Fab^H^ respectively. (**H**) Model of cell killing experiment. One Fab recognizes EGFR extracellular domain on the antigen presenting cells (APC), while the second Fab binds to CD3 of the T-cell receptor. (**I**) The effect of BEZ-IgG molecule on PBMC/PC3 (10:1) co-cultures. Constructs were added at 200 nM concentration and the effect of cell killing was measured using LDH release assay after 24 hours. As a control, all the individual components of the system (lane 1, 2 and 3) were tested and showed a marginal cell killing, while the robust effect was observed when both components of the system were present (lane 4). As a positive control, previously published plug-and-play BiTE technology was used (lane 5). Components of lanes: 1 GA1-Fc-GLM not loaded with Fabs; 2 GA1-Fc-GLM + UCHT1 Fab^LRT^; 3 GA1-Fc-GLM + EGFR Fab^H^; 4 GA1-Fc-GLM + UCHT1 Fab^LRT^ + EGFR Fab^H^; 5 plug-and-play BiTE (OKT3-GA1) + EGFR Fab^LRT^.

Next, we asked if we could generate a set of plug-and-play IgGs to facilitate the orthogonal pairs of GA1-Fab^LRT^ and GLM-Fab^H^, allowing for the co-binding detection of two antibodies (Figure 4B). Such technology would be beneficial for detecting low abundant cell surface targets, where the avidity effect could increase the detected signal. To do so, we linked GLM to a murine Fc fragment and GA1 to a human one, creating the set of reagents that could be detected using anti-Human Fc or anti-Murine Fc secondary antibody. We performed a similar co-binding experiment using the HCC1954 cell line and two model cell surface receptors, HER2 and EGFR. Anti-Her2 Fab was grafted into Fab^H^, while Fab^LRT^ scaffold was introduced into anti-EGFR Fab prior to mixing with GLM-Fc or GA1-Fc, respectively. The binding of anti-Her2 Fab^H^ + GLM-Fc and anti-EGFR Fab^LRT^ + GA1-Fc were detected using anti-murine-Fc-A647 or anti-human-Fc-A488, respectively. This resulted in the efficient detection of the simultaneous binding of EGFR and HER2 plug-and-play IgGs, which was recorded when co-staining was tested (Figure 4C). The controls of Isotype Fabs in each configuration did not produce detectable binding, proving the system’s efficiency (Figure 4D). We did not detect any exchange between the components of the system (Figure 4E, 4F).

### 3.4). Plug-and-Play bispecific antibodies (bi-IgG)

Further, using “knobs-into-holes” heterodimerization technology (Dixit et al., 2013), we generated a bispecific IgG-like assembly that provides for the orthogonal usage of Fab^LRT^ and Fab^H^ (Figure 4G). The body of the Fc unit comprises a heavy chain component containing a GLM arm that binds specifically to Fab^H^ and a second heavy chain component composed of a GA1 arm that binds Fab^LRT^. The specificities of the arms prevent any possible interchanging the Fabs. To test the efficiency of this bi-IgG assembly, we engineered a bispecific T-cell engager (BiTE) using the combination of the GLM and GA1 heavy chain components (Figure 4H). The bi-IgG forces the engagement of an antigen presenting cancer cell (APC) with a cytotoxic T-cell. The APC in this experiment was a PC3 cancer cell highly expressing EGFR on its cell surface; this cell was targeted by an anti-EGFR Fab^H^ binding through the GLM arm. The second arm consisted of the GA1 variant, which binds to an anti-CD3 Fab^LRT^ (UCHT1) (Figure 4G). This targets and activates the T-cell receptor on a T-cell. The efficiency of the bi-IgG BiTE was determined by measuring the activity of the cytoplasmic enzyme lactate dehydrogenase (LDH), which gets released into the medium upon cell killing. The addition of bi-IgG that contained all the active components to PBMC-PC3 co-cultures at 200 nM, resulted in a robust cell killing (Figure 4I). Importantly, no effect was observed when only one of the arms of bi-IgG was present in the assembly (Figure 4I). Notably, the functional readout was similar to the positive control that applied OKT3 antibody in the previously described plug-and-play bi-Fab format (Slezak et al. 2020).

### 3.5). Secondary reagents for antibody binding detection by microscopy and flow cytometry

Co-staining of multiple targets using traditional antibodies for flow cytometry or microscopy experiments involves a careful selection of secondary detection reagents, often requiring the primary antibodies to be from different species to avoid cross-reactivity. In this regard, the high specificity of the engineered PG variants and the ease to which fluorescent labels can be attached to them makes them good candidates for myriad cell biology applications. To evaluate their performance, it was first important to assess the detection sensitivity of the system. The model system we used was simple and controllable, consisting of a GA1-SNAP tag fusion labeled with Alexa 647. It was then complexed to a Fab that recognizes the unliganded state of maltose binding protein (MBP), which had been engineered to be expressed on the surface of HEK293 cells. The anti-MBP Fab (7O) was converted to the Fab^LRT^ framework, providing a 100 pM binding affinity to the GA1-A647 modality. This high affinity between GA1-Fab^LRT^ allows for simply premixing all reagents prior to the staining protocol, eliminating an additional washing step and significantly shortening the procedure time. This model system allows for additional experimental control since the conformational change of MBP upon maltose binding results in the modulating 7O binding in a controlled way. Thus, the detection of an Alexa 647 signal in a flow cytometry experiment can be evaluated in a concentration-dependent manner and completely eliminated by the spike of 1 mM maltose (Figure S7). To further extend the utility of the approach, we next evaluated the feasibility of attaching SNAP G-Fc to commercial antibodies. Since most of the community that routinely uses antibodies relies on commercially available IgGs, we posited that the SNAP G-Fc modality would be broadly useful by providing an off-the-shelf means to label virtually any antibody derived from multiple species. We used a commercially available Lamin A/C Rabbit IgG for the test case. SNAP G-Fc and SNAP GA1 were labeled with Alexa 488 and Alexa 647 via SNAP-tag, respectively and combined with their respective antibody format. Hela cells were stained with anti-EGFR Fab^LRT^ for 15 minutes on ice to prevent the internalization of EGFR. The cells were then washed, fixed and permeabilized, followed by the staining with anti-Lamin A/C IgG. The binding of anti-EGFR and anti-Lamin A/C was detected with SNAP GA1-A647 and SNAP G-Fc-A488. The signal for anti-EGFR Fab^LRT^ was nicely distributed in the cell plasma membrane while the Lamin A/C was concentrated around the nuclear envelope (Figure 5).

**Figure 5.**
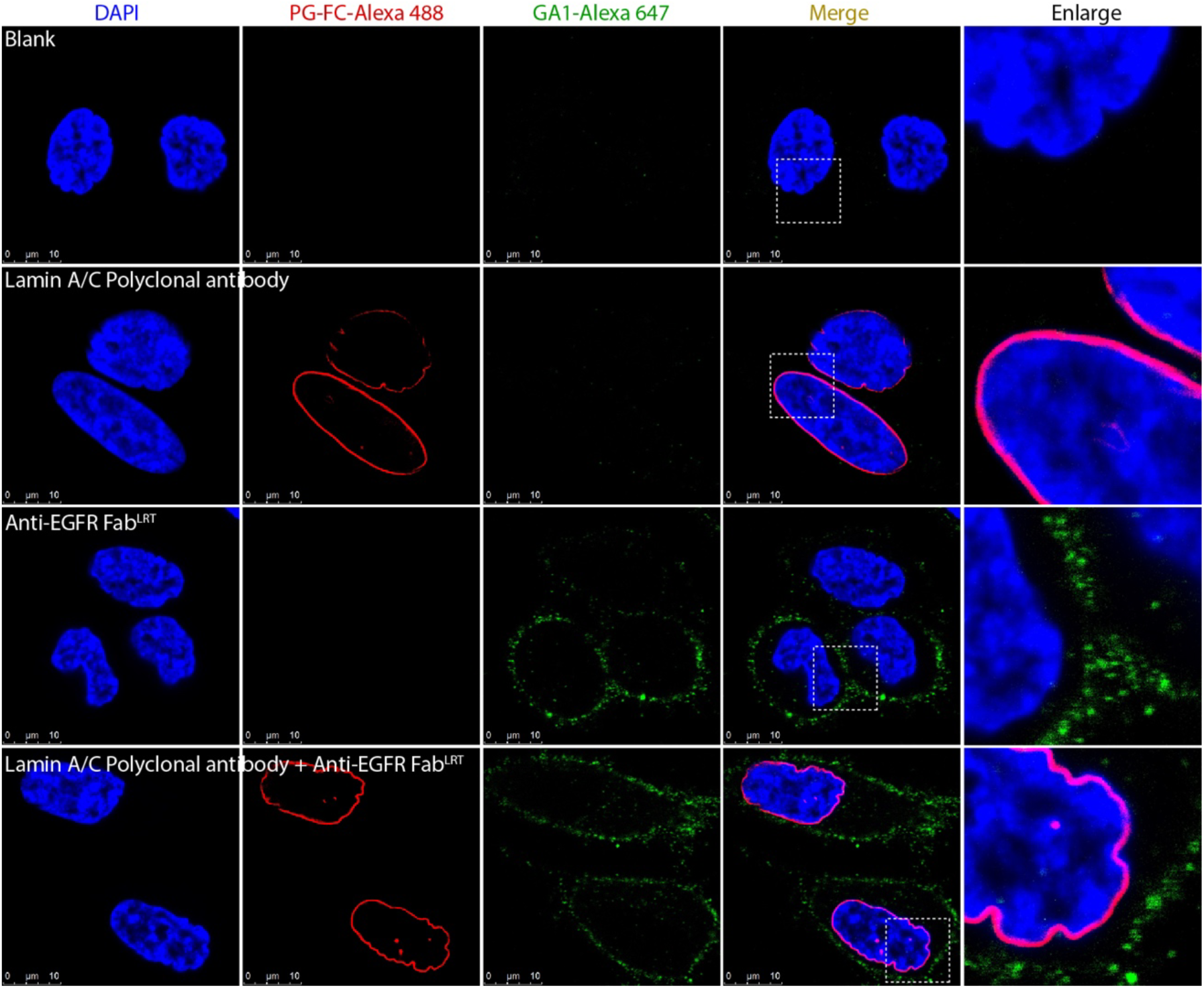
Engineered protein G allows for simultaneous co-staining using synthetic Fab^LRT^ and commercially available IgG. Synthetic Fab^LRT^ against EGFR and Lamin A/C Rabbit IgG were premixed with labeled protein GA1 and protein G-Fc and used for HeLa cell line staining. Immunofluorescence analysis shows the signal equally distributed signal for EGFR Fab^LRT^ in the plasma membrane while the Lamin A/C concentrated around the nuclear envelope. Orthogonal proteins GA1 and G-Fc shows a high specificity which allows for simultaneous co-detection.

Similar results were obtained for the co-binding detection of two fluorescently labeled Fabs that recognize different cell surface targets (Figure S9A). As a proof-of-concept, we chose to co-stain the HER2 and EGFR on SKBR3 cells, exploiting the ability to selectively match Fab scaffold types with their cognate PG variant partners. Before cell staining, SNAP GA1-A647 and SNAP GLM-A488 were premixed with anti-EGFR Fab^LRT^ and anti-HER2 Fab^H^ to form the two secondary detection agents. The binding of anti-EGFR and anti-HER2 was detected by SNAP GA1-A647 and SNAP GLM-A488 both independently and simultaneously (Figure S9B, C). Conversely, no co-staining of the cells was observed when the control isotype Fabs against Ebola nucleoprotein and MBP in analogous Fab-PG formats were used (Figure S9B). Not surprisingly, the fusion of double SNAP tag to GA1 significantly improved signal to noise ratio of the system (Figure S9D, S11).

## Discussion

We have designed and developed a versatile toolkit of powerful antibody-based reagents. These reagents can be assembled from independent functional component parts in a plug-and-play fashion using minimal molecular biology manipulation. This facilitates myriad experimental options that involve mixing and matching a few simple parts that can be combined to form multi-specific and multi-functional targeting moieties. These reagents are based on Protein G (PG)-Fab assemblies. These multifunctional modules can be constructed akin to Lego blocks, pieces of which can be prefabricated as standalone units and then linked together in a plug-and-play fashion. The ability to fabricate complex modular assemblies by snapping together the easy-to-manipulate component parts opens up the possibility of building highly diverse antibody-cargo combinations.

In previous work, we generated several high-affinity PG-Fab modules by modifying both PG and the Fab at their respective interaction surfaces using phage display (Bailey et al. 2014; Slezak et al. 2020). The resulting PG (GA1) and Fab (Fab^LRT^) variants resulted in an ultra-high affinity module facilitating cargo attachment at near covalent efficiency (100 pM) (Slezak et al. 2020). The toolkit described here complements and further extends the capabilities of this technology. Based on the initial success, we realized the plug-and-play concept could become even more powerful by developing a series of individual complementary Fab-PG pairs with differentiable targeting and functionality properties when linked together. Notably, the GA1 variant binds only to modified Fab scaffolds, not *wt* human or murine Fabs (Bailey et al. 2014). Thus, we undertook a phage display campaign to generate a new cohort of PG variants that were specific to several different *wt* Fab scaffolds, for instance, kappa vs. lambda light chains or human vs. murine scaffolds. This provided us with two categories of PG variants: one that binds only to modified Fab scaffolds (GA1) and several that were selective to different *wt* ones (GD, GF, and GLM).

We demonstrated that in several applications, these unique orthogonal pairs of PG and Fab modules can be combined as interchangeable parts, allowing a broad range of combinations to be evaluated in a multiplexed fashion. The constructs can be simple, like linking multiple copies of a PG variant like beads on a string to increase avidity to much more complex manifestations, which would link multiple different PG variants together to introduce muti-specificity into the assembly. Further, PG and its different variants are highly robust molecules to which various types of cargo can be readily fused (Slezak et al. 2020).

Among the most prevalent needs of cell biologists is the precise detection of specific antigens in complex cell environments. In this aspect, the PG-Fab modules can contribute significantly. This is done by fusing a SNAP-tag to the PG moiety and attaching it to a Fab targeting the antigen of interest. The SNAP-tag can be functionalized with a broad range of fluorophores, providing a spectrum of color options. Alternatively, GFP, YFP, or other fluorescent proteins can be fused to a PG variant to provide similar detection alternatives. These strategies achieve much improved signal-to-noise than traditional secondary polyclonal antibodies, eliminating extensive wash steps required to reduce background.

The development of orthogonal pairs of PG and Fab scaffolds that do not exchange binding partners creates an opportunity for the simultaneous detection of two different Fabs recognizing distinct antigens on a cell surface. Currently, immunostaining of multiple antigens involves combining secondary antibodies labeled with fluorescent probes possessing distinct emission maxima. These protocols require the use of antibodies from different species to avoid “cross-talk”, which often limits and complicates experimental design (Krenacs, Krenacs, and Raffeld 2010). To circumvent these problems, we labeled SNAP-fused GA1 (only binds modified Fab scaffolds) and GLM (only binds wt Fab scaffolds) with Alexa 647 and Alexa 488, respectively, to enable their simultaneous detection via flow cytometry. For proof-of-concept, we grafted anti-EGFR and anti-Her2 Fabs into engineered Fab^LRT^ and *wt* Fab^H^ scaffolds to detect their simultaneous binding with labeled GA1 and GLM (Figure S9). The experimental design is straightforward, involving a combined incubation with both Fabs premixed with their respective PG partners before the experiment. This resulted in efficient co-staining of anti-Her2 and anti-EGFR as detected on the target cells (Figure S9). Further, the PG variant detection moieties are off-the-shelf reagents facilitating antigen co-detection to be readily multiplexed to quickly test combinations of multiple pairs of targets using a plug-and-play format.

Importantly, similar concepts can be introduced into constructs that mimic full-length IgG formats. We designed an IgG-like construct containing two PG variants linked to either a human or murine Fc fragment. Depending upon the PG variant used, a Fab with a *wt* scaffold (Fab^H^ to GF) or an engineered one (Fab^LRT^ to GA1) can be attached to an Fc domain, either human or murine (Figure 4A). The PG-Fc IgG-like assemblies are highly stable and can be stockpiled to be used as off-the-shelf reagents. This provides the ability to go directly from Fab to IgG in minutes, not days or weeks. Additionally, bispecific PG-Fab moieties in both human and murine formats can be fashioned in a straightforward manner using the conventional “knobs into holes” heterodimerization strategy (Figure 4G). One advantage of the construct is that the purification of the bispecific PG-Fc is trivial. The bispecific module can be produced and stored and is ready to be used with multiple combinations of Fabs in applications involving multi-specific targeting reagents. For example, we demonstrated its versatility by building a bispecific T cell engager (BiTE), introducing a Fab (UCHT1) targeting CD3 of a T cell on one arm with the other arm loaded with a Fab targeting EGFR on the cancer cell. We show this BiTE produces potent cell killing in an LDH cell killing assay. Notably, even though the connection between the Fab and the PG-Fc is not covalent, we show that it is more than sufficient to perform experiments over long time intervals without loss of functionality (Figure S5).

We also sought to introduce into the plug-and-play platform the ability to exploit the huge catalog of antibodies available through commercial sources. This enhancement greatly expands the target space. Ironically, this involved reverse engineering the PG scaffold since its ability to bind to human or murine Fc domains had been engineered out during the development of the original Fab-specific GA1 scaffold (Bailey et al. 2014). The mutations that were introduced in GA1 to inhibit Fc binding were reverted back to their *wt* sequence and high affinity binding (Kd=4 nM) to Fc was restored (Figure 3). Since GA1 was originally engineered to bind only to an unnatural Fab scaffold (4D5), the resulting PG-Fc does not bind any *wt* Fab scaffold. In this form, PG-Fc can attach to a wide variety of molecular cargos efficiently, from commercial antibodies to other fabricated IgG molecules. A chain of several PG-Fc domains would enable linking multiple IgGs together into a single super avidity assembly.

The examples presented demonstrate the potential scope of experiments that could be enabled by the ability to snap together easy to manipulate component parts to build highly diverse antibody-cargo combinations. The ease of fabricating these modular assemblies also meets the criterion of “user-friendly”, where relatively low-level molecular biology expertise suffices to construct them. The amalgamation of a comparatively small number of different PG-Fab (IgG) pairings provides for combinatorial expansion of the types of assemblies produced from simple multi-valent and multi-specific starting building blocks. Importantly, these building blocks can be produced in bulk and stored to be used as “off the shelf” reagents in future experiments or to facilitate multiplexing across a cohort of antibodies in a single high throughput experiment.

## Acknowledgements

We extend our sincere gratitude to the members of the Kossiakoff Laboratory. We are indebted to the staff at the University of Chicago Comprehensive Cancer Center Sequencing Facility for their technical support and expertise. Furthermore, we wish to express our appreciation to the personnel at the Advanced Photon Source Beamline 24-ID at Argonne National Laboratory for their assistance in the acquisition of diffraction data. It should be noted that several figures in this manuscript incorporate illustrations generated using BioRender.com. This project was funded by NIH grants R01GM117372 (to A.A.K.).

## Material and methods

### Protein cloning, expression, and purification

The sequences of all used constructs are provided in Table S5. Protein Gs cloning strategy was previously described (Slezak et al. 2020). Briefly, the protein Gs were cloned using Sma1 site into pEKD40 with the Thrombin cleavable N-terminal SNAP-tag and C-terminal His-tag. Fabs. All Fab^LRT^ scaffold was grafted into Fab light chain at aa positions 123-127 (SQLKS −> LRT) using quick change site-directed mutagenesis. Protein Gs were expressed in BL21 (DE3) grown overnight in 2xYT medium in 20°C post induction with 1 mM IPTG at OD_600_= 0.6. Cells were sonicated in buffer A containing 50 mM Tris-HCl, pH 8.0, 150 mM NaCl and 10% glycerol. His-tagged protein was purified from the supernatant post centrifugation using Talon (TaKaRa) cobalt resin and eluted with 100 mM imidazole in buffer A.

Fabs and expressed in the periplasm of *E. coli* BL21 cells for 4 hours at 37°C post induction with 1 mM IPTG at OD_600_= 0.8-1. The cells were harvested by sonication in Protein G-wash buffer (Bailey et al. 2014) (50 mM Phosphate buffer, 500 mM NaCl, pH 7.4). After centrifugation the supernatant was applied on the protein GF affinity column. Proteins were eluted from the column with 0.1 M glycine, pH 2.6, and neutralized with 1M Tris-HCl, pH 8.5.

Fc fusion proteins were cloned into the pSCSTa vector containing a human or murine IgG construct. First, the vector was linearized via PCR designed to remove the CH1 portion. GA1 was amplified by PCR to contain a C-terminal linker (GGGGGGSGGGGSGGGGSSSGSS) and was then cloned into the N-terminal portion of the CH2-CH3 construct remaining in the open vector. GLM-hFc and GLM-mFc were generated by site-directed mutagenesis using GA1-hFc and GA1-mFc as templates. PG-Fc fusions were produced by transfection of Expi293 cells (ExpiFectamine, Gibco) according to manufacturer’s recommendation and were purified with protein A resin.

### Protein GF resin preparation

Protein GF resin was generated as previously described (Bailey et al. 2014). Briefly, protein GF was cloned with a SUMO-tag, that contains a free cysteine to allow for covalent linkage to SulfoLink Coupling Resin (Thermo Scientific), and the resin was created following the manufacturer’s protocol.

### Phage display library preparation

The library generation strategy was designed using previously described approach (Sidhu et al. 2000; Bailey et al. 2014). Protein G was displayed on the surface of M13 phage by fusion to the minor coat protein pIII. After the inspection of the Fab-protein G crystal structures (1IGC and 6U8C) the 6 amino acids at the position 38-43 were identified to interact with the Fab constant light chain. These residues were randomized using hard randomization strategy (NNK) where all amino acids are possible. Stop codon was placed using quick change site-directed mutagenesis in the aa position 40 prior to ssDNA preparation from phage. Phosphorylated primers were used in Kunkel mutagenesis (Kunkel 1985) and the library was created as described before (Slezak and Kossiakoff 2021).

### Phage display selection

In the first round of selection, 1 μ*M* of Fab^H^ (Human Kappa) was immobilized on 200 μl SA magnetic beads (Promega) and incubated with 1 mL phage library for 1 hour at room temperature with gentle shaking. Each of the generated phage libraries were used separately. The beads were washed three times to remove nonspecific phage and added to log phase *E. coli* XL-1 blue cells (Stratagene) and incubated for 20 minutes in room temperature. Then, media containing 100 μg/mL ampicillin and 10^9^ p.f.u./mL of M13K07 helper phage (NEB) was added for overnight phage amplification in 37°C. For all subsequent rounds, the amplified phage was precipitated in 20% PEG/2.5 M NaCl for 20 minutes on ice. Before each round, phage pool was negatively selected against empty paramagnetic beads for 30 minutes with shaking to eliminate nonspecific binders. The final concentration of antigen was dropped gradually from 1 μ*M* to 1 nM from the first to the fifth round (2^nd^ round: 200 nM, 3^rd^ round: 50 nM, 4^th^ round 10 nM and 5^th^ round 1 nM). After phage binding, the beads were subjected to five washing rounds. The bound phages were eluted using 0.1 M glycine, pH 2.6 and neutralized with TRIS-HCl, pH 8.5. Then, the phage eluate was used for *E. coli* infection and phage amplification as described above. After round 4^th^ and 5^th^ phages were plated on ampicillin plates and 96 single colonies were picked for single-point phage ELISA assays. The promising clones demonstrating high ELISA signal and low non-specific binding were sequenced and reformatted into a pEKD40 expressing vector as described in Protein expression and purification.

To generate the universally binding protein G, the different fab scaffolds were introduced as a target in consecutive rounds of selection (1^st^ round: Fab^H^, 2^nd^ round: Fab^L^ (Human Lambda), 3^rd^ round: Fab^LRT^, 4^th^ round: Fab^S^ (4D5), 5^th^ round: Fab^H^). The selection procedure was then followed us described above.

### Enzyme-Linked Immunoabsorbent Assays (ELISA)

ELISA protocol was described before in (Lan et al. 2024). Briefly, 50 nM of a different Fab scaffolds were directly immobilized on high binding experimental wells (Greiner Bio) and BSA was immobilized in control wells, followed by extensive blocking with BSA. After 15 minutes incubation with phage, well were extensively washed three times and incubated with Protein L-HRP (Thermo Scientific, 1:5000 dilution in HBST) for 20 minutes. The plates were again washed and developed with TMB substrate (Thermo Scientific) and quenched with 10% H_3_PO_4_, followed by the absorbance at A_450_ determination.

### Multipoint ELISA

High-binding experimental wells (Greiner Bio) were used to immobilize the target proteins at a concentration of 50 nM. The wells were extensively blocked with BSA. Twelve 2-fold serial dilutions were added and incubated for 20 minutes at room temperature for each construct being analyzed. The wells were then subjected to extensive washing before being incubated with HRP-conjugated anti-human (Fab)2 antibody (JacksonImmunoResearch) at a dilution of 1:5000 in PBST for 20 minutes at room temperature. After washing again, the wells were developed with TMB substrate (Thermo Scientific), and the reaction was quenched with 10% H3PO4. The absorbance at A450 was then determined.

### Surface plasmon resonance analysis

All Surface plasmon resonance (SPR) analysis were performed on a MASS-1 (Bruker). All targets were immobilized via a 6x His-tag to a Ni-NTA sensor chip. Fabs in twofold dilutions were run as analytes at 30 μl/min flow rate at 20°C. Sensogram were corrected through double referencing and 1:1 binding model fit was done using Sierra Analyser (Bruker). To ensure the binding properties the experiments were repeated in an opposite order, where the His-tagged Fabs were immobilized to a Ni-NTA sensor chip and the Protein Gs were run as analytes.

### Fab^H^-GF and GD crystallization and structure determination

Recombinant Fab^H^ E12, its target ASF1 (Bailey et al. 2018) and protein G were produced and purified as described above. SNAP-tag was removed from the protein G by Thrombin cleavage in room temperature overnight and purified by IMAC on a Talon resin (TaKaRa). To obtain the ASF1-Fab^H^-protein GF or GD, the proteins were incubated in 1:1 molar ratio and the complex was purified on a size exclusion chromatography on a Sephadex 200 column, equilibrated with HBS. The purity of the complex was confirmed by SDS-PAGE. Both protein complexes were concentrated to 10mg/ml prior to initial crystallization trials set up at room temperature using the hanging-drop vapor-diffusion method utilizing the Mosquito Crystal robot (TTP Labtech).

The ASF1-Fab^H^E12-GF complex was crystallized by mixing 100nl of protein complex solution with 100nl of a Protein Complex Suite (NeXtal) screen solution. The most promising crystals were observed in 0.1M Sodium Cacodylate pH 5.5, 0.1M Calcium acetate hydrate and 12% PEG 8000. Crystal quality was improved by the condition optimization and seeding. Hanging-drop crystallization was set up by mixing 1 μl of a complex with 1 μl of reservoir solution. Quality was further improved by the seeding technique (Luft and DeTitta 1999) in 0.1M Sodium Cacodylate pH 6.0, 0.1M Calcium acetate hydrate and 10% PEG 8000 at room temperature. The crystals were soaked in mother liquid containing 20% PEG 400 as a cryoprotectant and flash-frozen in liquid nitrogen for data collection.

The ASF1-Fab^H^E12-GD complex was crystallized by mixing 100nl of protein complex solution with 100nl of a JCSG Top96 (Rigaku) screen solution. The most promising crystals were observed in 0.1M Sodium cacodylate pH 6.5 and 1M Sodium citrate tribasic. Crystal quality was improved by the condition optimization and seeding. Hanging-drop crystallization was set up by mixing 1 μl of a complex with 1 μl of reservoir solution. Quality was further improved by the seeding technique (Luft and DeTitta 1999) in 0.1M Sodium cacodylate pH 6.8 and 0.8M Sodium citrate tribasic at room temperature. The crystals were soaked in mother liquid containing 20% PEG 400 as a cryoprotectant and flash-frozen in liquid nitrogen for data collection.

X-ray diffraction datasets were collected at the NECAT 24-ID-E beamline at the Advanced Photon Source. Crystal structures were determined by molecular replacement method using the structures of the Fab-ASF1 complex (pdb: 6AYZ) and protein G (pdb: 6U8C) using PHASER (McCoy et al. 2007). The structure refinements were done using phenix.refine software (Afonine et al. 2012). The electron density maps and the manual corrections were performed using COOT (Emsley and Cowtan 2004). Structural figures were created using ChimeraX (Goddard et al. 2018). Coordinates have been deposited to Protein Data Bank under entry: 9AWE and 9AVO.

### Flow cytometry

SNAP-GA1 and SNAP-GLM were conjugated with BG-Alexa Fluor 647 and BG-Alexa Fluor 488 (New England Biolabs) respectively, according to the manufacturer’s protocol. Excess substrate was removed with PD MiniTrap G-25 (Cytiva) desalting columns.

A human embryonic kidney (HEK) cell line that was stably transfected to express extracellular MBP anchored to a transmembrane domain and intracellular green fluorescent protein (HEK^M-TM-G^) was used for initial assessment of SNAP-GA1-A647 as a secondary detection reagent. HEK^M-TM-G^ cells were cultured to ~70% confluency before detachment by trypsin digestion. Cells were washed once in PBS/1% bovine serum albumin (BSA) and added at a concentration of 500,000 cells per tube to Eppendorf tubes. Cells were incubated with no Fab, 7O^LRT^, 7O^LRT^ in 10 mM maltose, and MJ6^LRT^ for 30 minutes at room temperature before washing two times in 1 mL PBS/1% BSA. Next, Alexa Fluor 647 AffiniPure Goat Anti-Human IgG F(ab’)_2_ fragment specific (Jackson ImmunoResearch) and SNAP-GA1-A647 were added to samples for 20 minutes at room temperature. Cells were resuspended in 250 uL PBS/1% BSA after the final wash and subjected to analysis by a CytoFLEX flow cytometer (Beckman Coulter).

The human breast cancer cell line SKBR3 (ATCC) was cultured to ~70% confluency before detachment by trypsin digestion. Cells were washed once in PBS/1% bovine serum albumin (BSA) and added at a concentration of 500,000 cells per well to a 96-well round bottom plate. SNAP-GLM-A488 and SNAP-GA1-A647 were incubated at equimolar concentrations with combinations of MJ20^H^, HER2^H^, 7O^LRT^, and EGFR^LRT^ for 1 hour on ice before diluting to a final concentration of 250 nM for each component in PBS/1% BSA. Samples were incubated on cells for 30 minutes at room temperature before washing three times with 250 uL of PBS/1% BSA by centrifugation at 400 xg for 5 minutes. Cells were resuspended in 250 uL PBS/1% BSA after the final wash and subjected to analysis by a CytoFLEX flow cytometer (Beckman Coulter).

### Co-staining detection using flow cytometry

Co-staining detection of GA1-hFc and GLM-mFc with HER2^H^ and EGFR^LRT^ Fabs, respectively, was performed with the HCC1954 cell line. PG-Fc fusions were pre-incubated with HER2^H^, EGFR^LRT^, MJ20^H^ (isotype control), or S1^LRT^ (isotype control) on ice for 30 minutes. The EGFR^LRT^ + GA1-hFc assembly was first added to cells at 20 nM for 15 minutes on ice. Cells were washed three times with PBS/1% BSA before adding HER2^H^ + GLM-mFc at 20 nM for 15 minutes on ice. Cells were washed three times before adding a mixture of Alexa Fluor 488-conjugated Anti-Human Fc and Alexa Fluor 647-conjugated Anti-Mouse Fc for 15 minutes on ice followed by three washes.

### Co-staining detection using microscopy

Hela cells (ATCC) were trypsinized and resuspended in DMEM medium (Gibco) including 10% FBS, and then seeded in a cell culture chamber slide (Ibidi) at a density of 1.8X10^4^ cells/well following incubation in 37°C/5% CO_2_ for 24 h. Cells was treated with 100 nM anti-EGFR Fab^LRT^ in culture medium for 15 min in 37°C/5% CO_2_, and incubated with 100 nM GA1-Alexa 647 in 4°C for 1 h after being washed by PBS. Then cells were sequentially fixed by 4% PFA/PBS (Thermo Fisher) at RT for 15 min, permeabilized by 0.2%Triton X-100/PBS (Thermo Fisher) at RT for 20 min and blocked by 3% BSA/PBS at RT for 1 h. Lamin A/C Polyclonal antibody was diluted in 100 nM by 0.5% BSA/PBS and added to cells (150 μL/well) prior to incubation in 4°C overnight. Cells were washed and incubated with 100 nM PG-Fc-Alexa 488 in 4°C for 1 h. After being stained by DAPI (Thermo Fisher), cells were examined by confocal microscope (Leica).

### Avidity effect testing by the SPR

A multivalent form of protein GA1 was created by the introduction of a Gly-Ser linker. Protein GA1 was multimerized from dimer up to decamer with the spectrum of different linkers. The proteins were expressed and purified as described above. His-tagged Fab in Fab^s^ scaffold was immobilized on the Ni-NTA sensor chip. Monomer, dimer, and trimer GA1 in two-fold dilutions starting at 200 nM were run as analytes at 30 μl/min flow rate at 20°C. Sensogram were corrected through double referencing and 1:1 binding model fit was done using Sierra Analyser (Bruker).

### T-cell redirection cell-culture assay

Human prostate cancer cell line PC-3 (ATCC), overexpressing EGFR on the cell surface was cultured according to ATCC protocols. CD3-positive PBMC cells were isolated from blood (Vissers, Jester, and Fantone 1988). The day before the experiment, PC-3 cells were seeded into a 96-well plate (20K PC-3 cells in 100 μ*l* per well). The next day, PBMC cells were added to medium aspirated PC-3 cells at 10:1 Effector cell to Target cell ratio and then the bi-specific components were added at 200 nM. After 24 hours of co-culturing, the medium was analyzed for LDH presence using commercially available kit (CytoTox96, Promega). The results were analyzed and normalized using protocols and standards provided in the kit.

### Figures generations

The models in figures were generated using Biorender.com.

