## Supplementary materials for "Engineered Protein-G variants for plug-and-play applications"

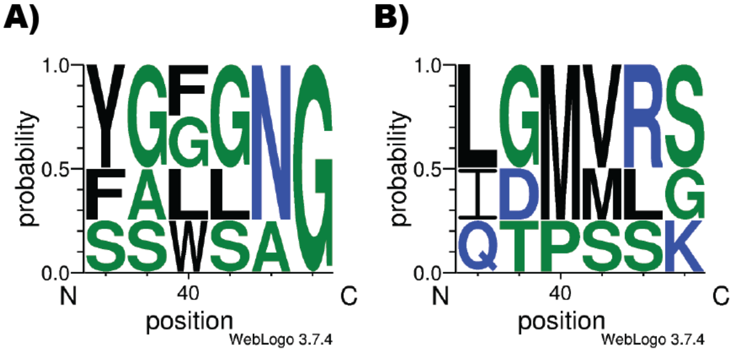


**Figure S1.** (**A**) Weblogo represents amino acids sequences of newly developed protein Gs that pose universal binding properties (binds all existing Fab scaffolds). (**B**) Weblogo represents protein Gs selectively binding only $Fab^{H}$ and $Fab^{L}$, but do not bind to $Fab^{LRT}$.

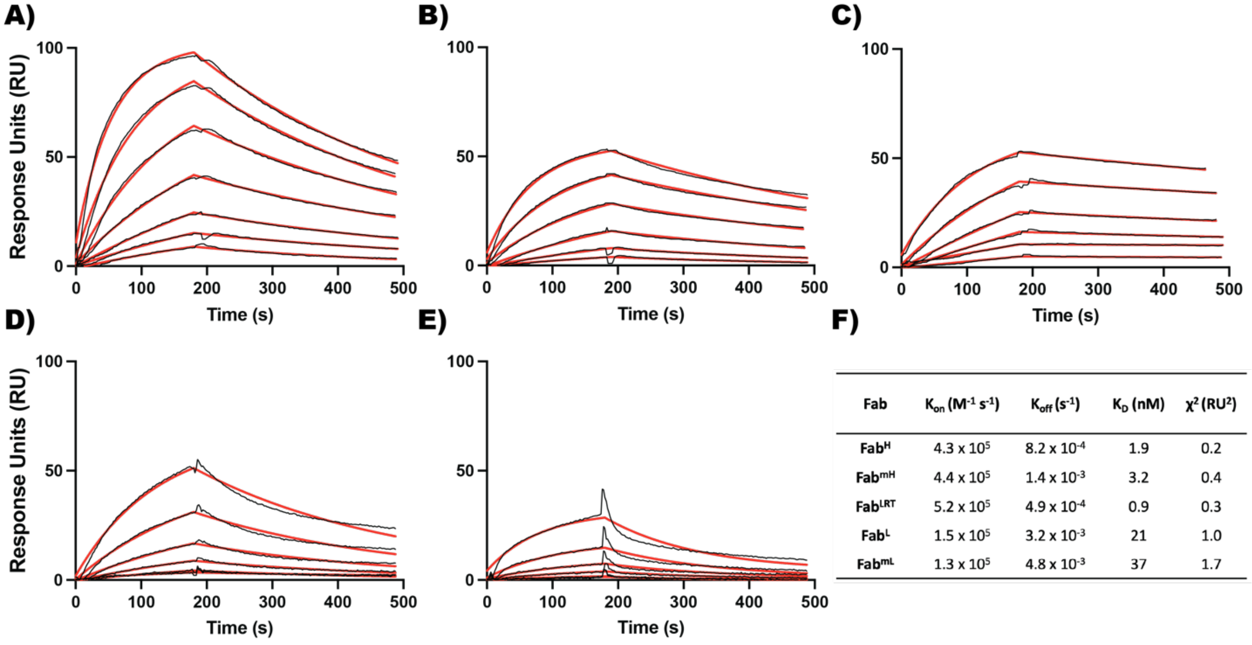


| Fab | $K_{on}$ ($M^{-1}$ $s^{-1}$) | $K_{off}$ ($s^{-1}$) | $K_D$ (nM) | $\chi^2$ ($RU^2$) |
|---|---|---|---|---|
| $Fab^{H}$ | $4.3 \times 10^{5}$ | $8.2 \times 10^{-4}$ | 1.9 | 0.2 |
| $Fab^{mH}$ | $4.4 \times 10^{5}$ | $1.4 \times 10^{-3}$ | 3.2 | 0.4 |
| $Fab^{LRT}$ | $5.2 \times 10^{5}$ | $4.9 \times 10^{-4}$ | 0.9 | 0.3 |
| $Fab^{L}$ | $1.5 \times 10^{5}$ | $3.2 \times 10^{-3}$ | 21 | 1.0 |
| $Fab^{mL}$ | $1.3 \times 10^{5}$ | $4.8 \times 10^{-3}$ | 37 | 1.7 |

**Figure S2. Characterization of GF- an universal Fab binder.** (A) SPR sensogram showing the interaction with $Fab^{H}$. (B) SPR sensogram showing the intraction with $Fab^{mH}$. (C) SPR sensogram showing the interaction with $Fab^{LRT}$. (D) SPR sensogram showing the interaction with $Fab^{L}$. (E) SPR sensogram showing the interaction with $Fab^{mL}$. (F) Kinetic binding parameters. For the kinetic experiment, fabs were serially diluted two-fold, starting at 50 nM for all Fabs, and 25 nM for $Fab^{LRT}$.

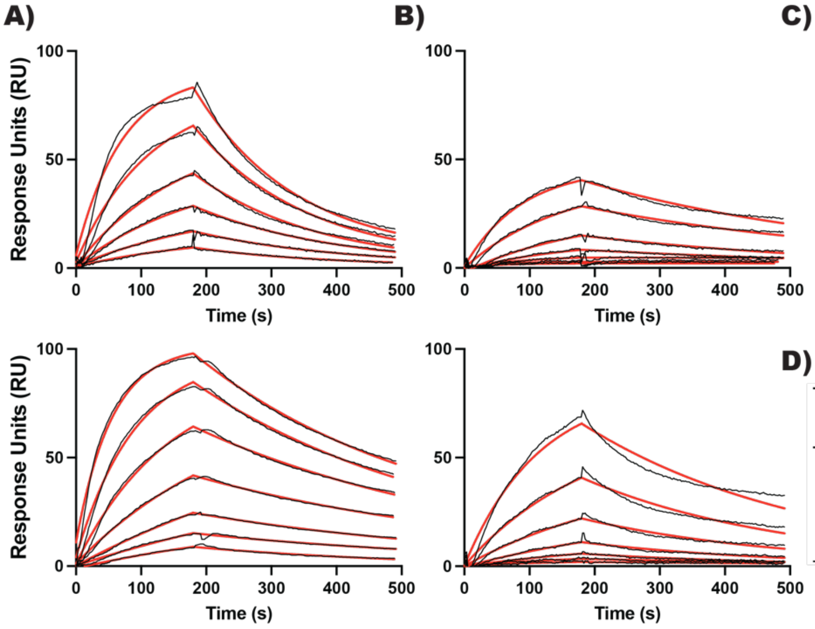


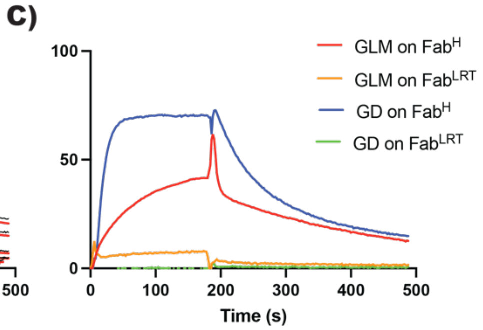


D)

| pG | Fab | $K_{on}$ ($M^{-1}$ $s^{-1}$) | $K_{off}$ ($s^{-1}$) | $K_D$ (nM) | $\chi^2$ ($RU^2$) |
|---|---|---|---|---|---|
| D | $Fab^H$ | $8.3 \times 10^5$ | $5.3 \times 10^{-3}$ | 6.4 | 1.3 |
| | $Fab^L$ | $1.3 \times 10^5$ | $2.1 \times 10^{-3}$ | 16 | 0.7 |
| LM | $Fab^H$ | $2.6 \times 10^5$ | $2.2 \times 10^{-3}$ | 8.9 | 0.7 |
| | $Fab^L$ | $1.4 \times 10^5$ | $3.0 \times 10^{-3}$ | 21 | 2.1 |

**Figure S3. Characterization of GD and GLM** (A) SPR sensogram showing the interaction of GD (top) and GLM (bottom) with $Fab^H$. (B) SPR sensogram showing the interaction of GD (top) and GLM (bottom) with $Fab^L$. (C) A single injection of $Fab^H$ and $Fab^{LRT}$. 25nM of each fab was injected, and no binding to $Fab^{LRT}$ was observed. (D) Kinetic parameters of binding. For the kinetic experiment, Fabs were serially diluted two-fold, starting at 100nM.

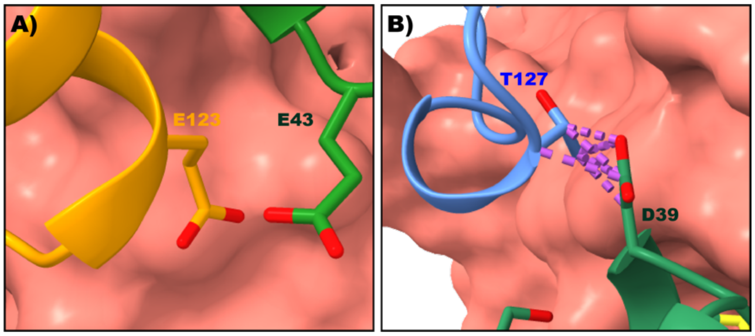


**Figure S4 Structural basis of an orthogonal specificity between GA1-$Fab^{LRT}$ and GD-$Fab^H$.** (A) Model of GA1-$Fab^H$. The charge clash of E123 from $Fab^H$ and E43 from GA1 is responsible for no interaction between these molecules. (B) Model of GD-$Fab^{LRT}$. Light chain loop rearrangement caused by two amino acids deletion in $Fab^{LRT}$ causes a clash between T127 and D39 from the GD. Molecules are colored as follows: Fab Hc- red, $Fab^H$ Lc- orange, $Fab^{LRT}$ Lc- blue, GA1, and GD- green.

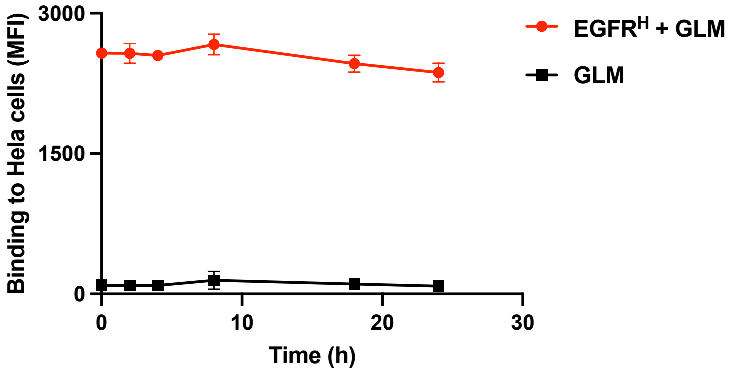


**Figure S5**. **Stability of Plug and Play PG complexes in cell-based applications.**

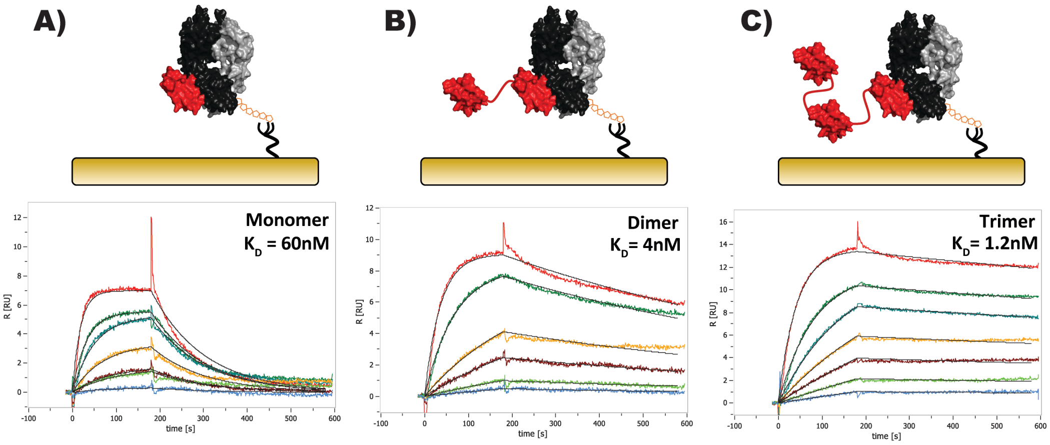


**Figure S6. Avidity-driven GA1 dissociation rate enhancement.** (A) SPR sensogram showing the interaction with pG-A1 monomer and Fab$^S$. (B) SPR sensogram showing the interaction with pG-A1 dimer and Fab$^S$. (C) SPR sensogram showing the interaction with pG-A1 trimer and Fab$^S$. The kinetic parameters exhibit substantially decreased $k_{off}$ values with increasing pG-A1 valency but only marginally decreased $k_{on}$ values.

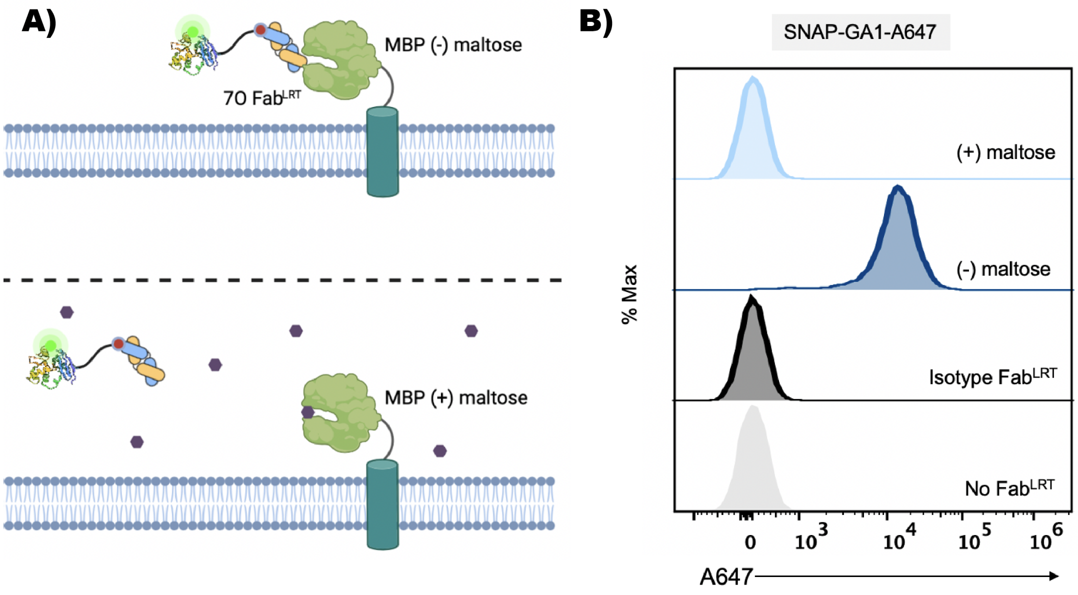


**Figure S7. Labeled SNAP-protein G as a tool for antibody binding detection by flow cytometry**. (**A**) Experimental model. A conformation specific anti-MBP 7O Fab$^{LRT}$ was used to detect the extracellular MBP stably engineered on the surface of the HEK cell line. Conformational change of MBP upon maltose addition eliminates the anti-MBP 7O Fab$^{LRT}$ binding. SNAP-GA1 labeled with Alexa 647 is used for the detection in flow cytometry and is premixed with fab before the cell staining. (**B**) Flow cytometry histogram of anti-MBP 7O Fab$^{LRT}$ binding to HEK cell line stably expressing extracellular MBP without the presence of the maltose. The system exhibits a strong signal with a shallow background. Fab binding is abolished upon maltose addition. As a negative control the SNAP-GA1 alone and the isotype fab against Ebola nucleoprotein were used. No detectable signal was observed. Fab is premixed with SNAP-GA1 before cell staining, which significantly shortens the staining protocol.

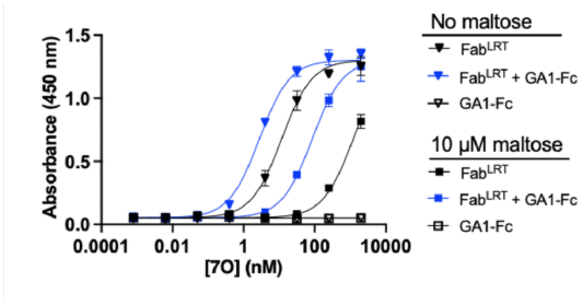


| | 7O sAB $EC_{50}$ [nM] | |
|---|---|---|
| Construct | No maltose | 10 µM maltose |
| Fab$^{LRT}$ only | 10.2 | >500 |
| Fab$^{LRT}$ + GA1-Fc | 2.97 | 85.1 |

**Figure S8. GA1-Fc fusion enables modular assembly of bivalent IgG-like sABs.** (**A**) ELISA $EC_{50}$ analysis of sAB 7O binding to immobilized MBP in the absence of maltose or with 10 µM maltose. In both cases, the IgG-like sAB format improves the 7O $EC_{50}$ value substantially. (**B**) Table of 7O $EC_{50}$ values.

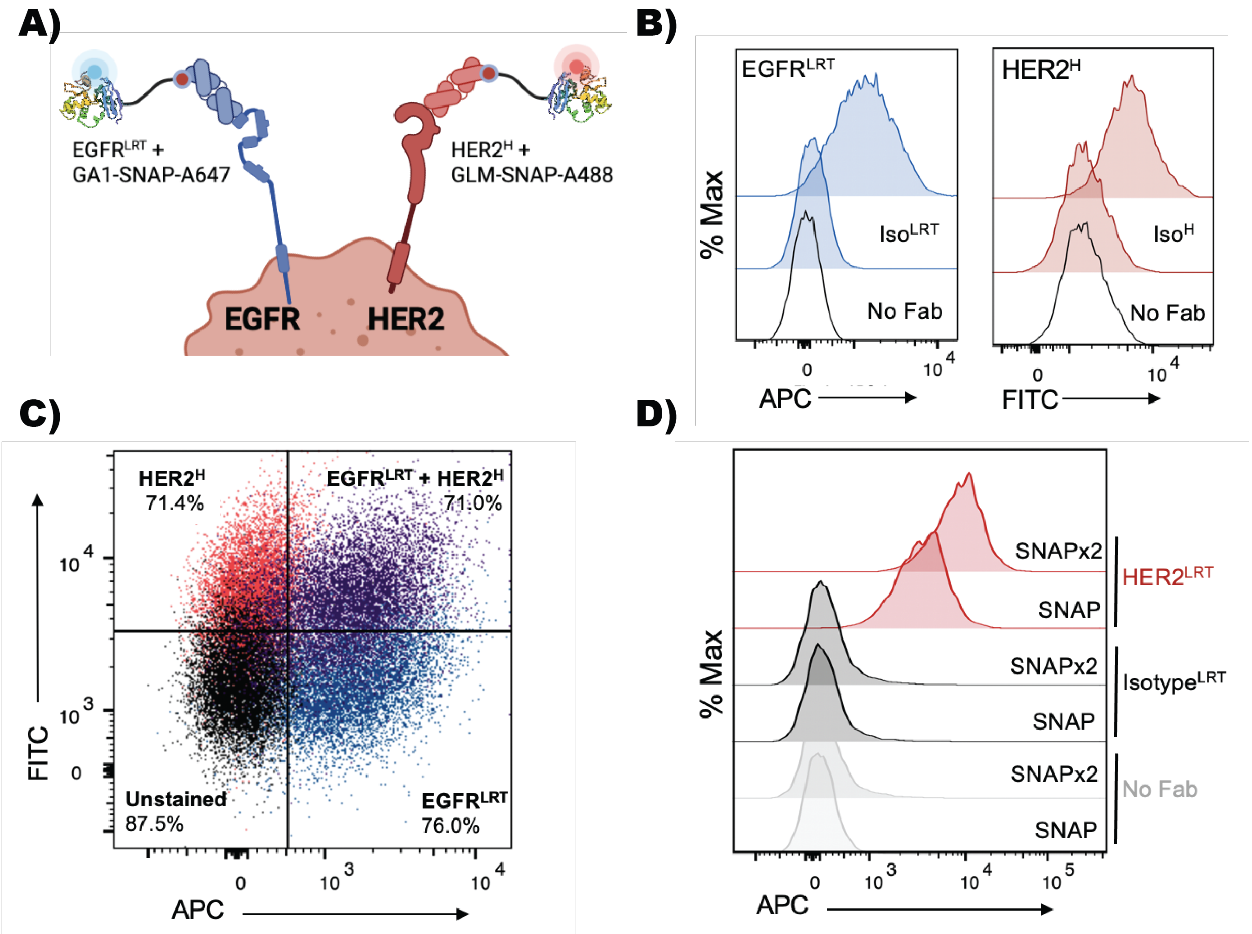


**Figure S9. Flow cytometry simultaneous binding detection of two different Fabs by the orthogonal pairing of GA1-Fab$^{LRT}$ and GLM-Fab$^{H}$. (A)** Model of secondary co-detection using the orthogonal pairing of GA1-Fab$^{LRT}$ and GLM-Fab$^{H}$. SNAP-GA1 and SNAP-GLM fusions are labeled with benzylguanine- (BG) Alexa 647 and Alexa 488, respectively, and then incubated with Fab$^{LRT}$ and Fab$^{H}$ molecules before addition to cells. Protein GA1 and GLM scaffold specificity allow for the simultaneous detection of the other fab binding on the cell surface. **(B)** Flow cytometry analysis of SKBR3 cell surface receptors. EGFR was detected by an anti-EGFR Fab$^{LRT}$ using SNAP-GA1-A647 as a secondary detection agent (*left*). HER2 was detected by an anti-HER2 Fab$^{H}$ using SNAP-GLM-A488 as a secondary detection agent (*right*). **(C)** Density plot analysis of **s**imultaneous detection of anti-EGFR Fab$^{LRT}$ and anti-HER2 Fab$^{H}$ via flow cytometry using SNAP-GA1-A647 and SNAP-GLM-A488, respectively, as secondary detection agents. The high degree of specificity for each GA1-Fab$^{LRT}$and GLM-Fab$^{H}$ interactions allows all Fab and secondary detection components to be mixed in one tube before adding to cells. The low background and depleted off-target recognition are demonstrated using Fab$^{LRT}$ and Fab$^{H}$ isotype controls. **(D)** Double SNAP fusion significantly increased detected signal.

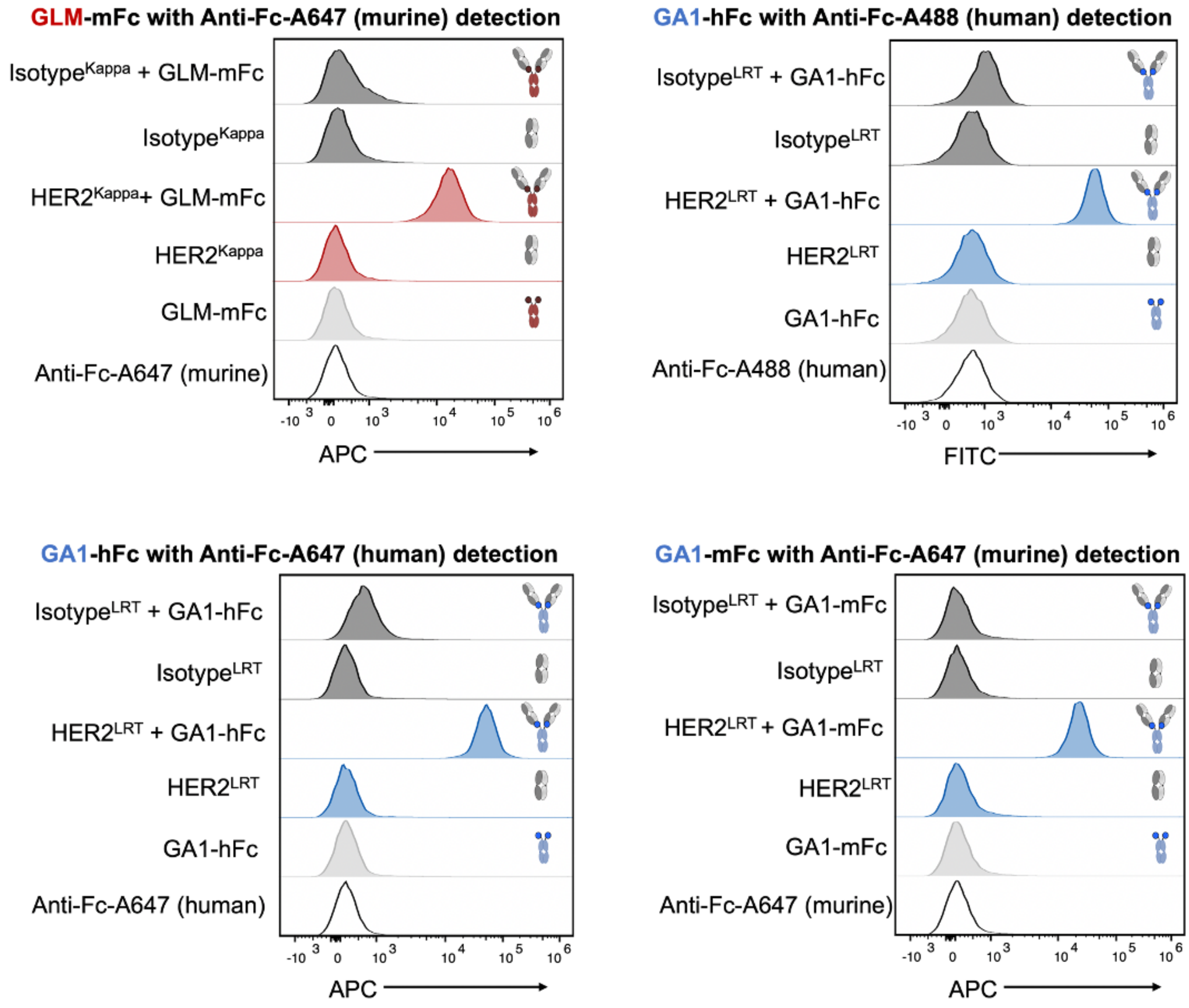


**Figure S10. Cell surface staining controls for the modular assembly of protein G-Fc fusions with HER2 targeting sABs.** MDA-MB-453 cells were stained with IgG-like assembled sABs targeting EGFR (Fab$^{LRT}$ format) or HER2 (Fab$^{K}$ format) at 200 nM. No background signal was detected using Fc-fusion only, Fab only, or the isotype controls.

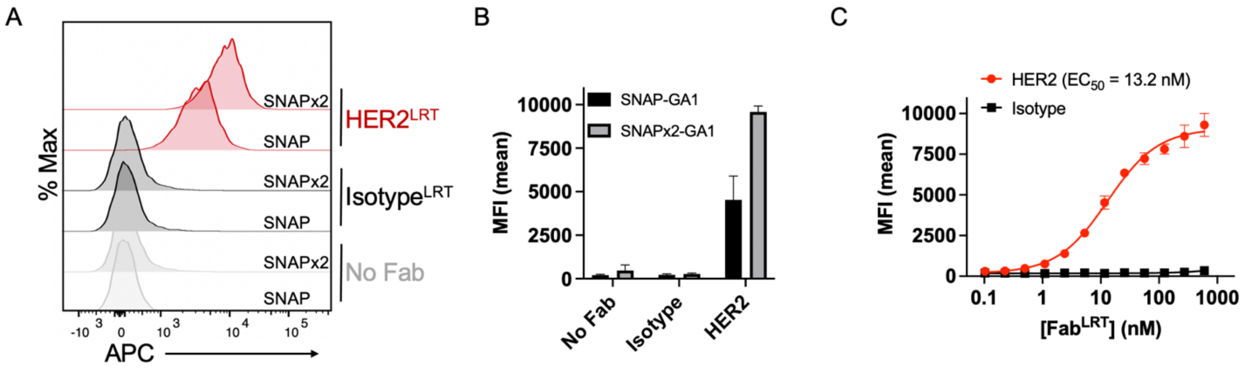

**Figure S11. GA1 fusion to tandem SNAP-tag enables low wash cell staining protocols.** A) MDA-MB-453 cells were stained with HER2 targeting Fab$^{LRT}$ or isotype control Fab$^{LRT}$ at 200 nM. Fabs were pre-incubated at an equimolar ratio with SNAP-GA1 or SNAPx2-GA1 labeled with Alexa Fluor 647. B) MFI quantification of the results in A shows that two SNAP-tags labeled with Alexa Fluor 647 doubles the signal. C) Cell surface $EC_{50}$ using 50 nM SNAPx2-GA1 labeled with Alexa Fluor 647 as a secondary detection agent.

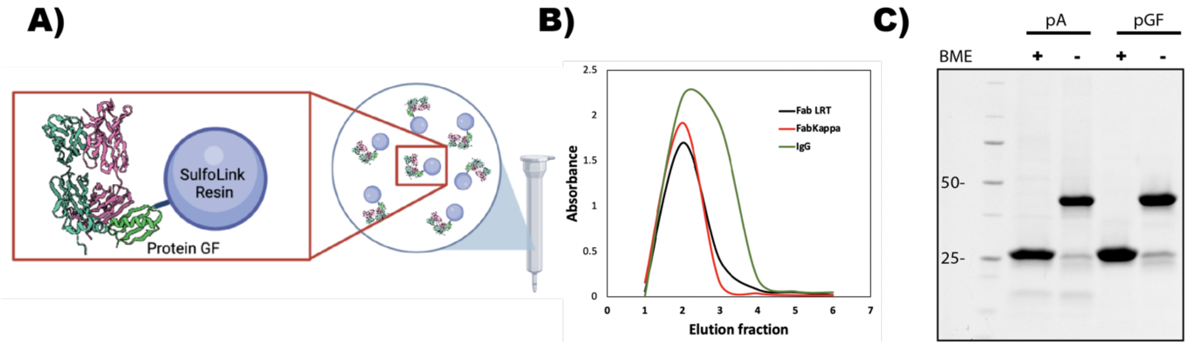


**Figure S12. In-house generation of affinity chromatography resin for universal Fab purification.** (A) Schematic showing the covalent immobilization of SUMO-GF with SulfoLink Coupling Resin. The resin is a high capacity (20mg of Fab/mL), has a long lifespan (>1.5 years), and is reusable due to its tolerance to low pH regeneration. (B) Elution profiles of full-length IgG, Fab$^{H}$, and Fab$^{LRT}$ purified by a single step with a SUMO-GF coupled resin. (C) SDS-PAGE gel post purification. Purity of the product is superior to the commonly used Protein-A resin due to its single chain binding profile.

| | Protein GD | Protein GF |
|---|---|---|
| Wavelength (Å) | 0.9791 | 0.9791 |
| Source | APS 24-ID-E | APS 24-ID-C |
| Resolution (Å) | 3 | 2.8 |
| Space group | I 41 2 2 | P 2 21 21 |
| Cell parameters (Å) | 134.97, 134.97, 339.57 | 57.87, 85.66, 180.11 |
| ( ° ) | 90.0, 90.0, 90.0 | 90.0, 90.0, 90.0 |
| Total reflections | 397,629 | |
| Unique reflections | 31,838 | 21810 |
| Multiplicity | 12.5 (13) | 1.9 (1.9) |
| Completeness (%) | 99.8 (100) | 95.6 (97.38) |
| Mean I/sigma(I) | 12.9 (2.1) | 7.7 (1.7) |
| R-merge | 0.16 (1.60) | 0.048 (0.444) |
| R-pim | 0.07 (0.65) | 0.048 (0.444) |
| CC1/2 | 0.99 (0.88) | 0.99 (0.70) |
| Reflections used in refineme | 31,735 | 21781 |
| R-work/R-free | 0.22/0.24 | 0.24/0.28 |
| RMS bond length (Å) / angle | 0.009/1.2 | 0.01/1.4 |
| Ramachandran favored (%) | 93.7 | 93.26 |
| allowed (%) | 4.42 | 5.46 |
| outliers (%) | 1.89 | 1.28 |

*values in parentheses are for highest-resolution shell.

**Table S1. Data collection and refinement statistics summary**

| Protein G | Amino acid sequence in position 38-43 | $K_{on}$ ($M^{-1}$ $s^{-1}$) | $K_{off}$ ($s^{-1}$) | $K_D$ (nM) |
|---|---|---|---|---|
| GF | YAFGNG | $4.3 \times 10^5$ | $8.2 \times 10^{-4}$ | 1.9 |
| GD | IDMVSS | $8.3 \times 10^5$ | $5.3 \times 10^{-3}$ | 6.4 |
| GLM | LGMMRS | $2.6 \times 10^5$ | $2.2 \times 10^{-3}$ | 8.9 |
| GS | SGLLAG | $2.8 \times 10^5$ | $1.6 \times 10^{-3}$ | 5.8 |

| | | | | |
|---|---|---|---|---|
| GLV | LGMVRG | $2.9 \times 10^{5}$ | $4.8 \times 10^{-3}$ | 16.5 |
| GYG | YGTANG | $3.6 \times 10^{5}$ | $2.6 \times 10^{-3}$ | 7.2 |
| GQT | QTPSLK | $2.5 \times 10^{5}$ | $3.2 \times 10^{-3}$ | 13.0 |
| GW | FGWSNG | $2.7 \times 10^{5}$ | $1.2 \times 10^{-3}$ | 4.3 |
| GY | YSGGNG | $3.7 \times 10^{5}$ | $1.4 \times 10^{-3}$ | 3.9 |
| GFH | FAHGNA | $2.0 \times 10^{5}$ | $2.1 \times 10^{-3}$ | 10.4 |
| GFS | FGNSNG | $2.3 \times 10^{5}$ | $1.9 \times 10^{-3}$ | 8.3 |

**Table S2. Kinetic parameters of different Protein GS binding to Fab$^{H}$.**

| Protein G | Amino acid sequence in position 38-43 | $K_{on}$ ($M^{-1}$ $s^{-1}$) | $K_{off}$ ($s^{-1}$) | $K_D$ (nM) |
|---|---|---|---|---|
| GF | YAFGNG | $5.2 \times 10^{5}$ | $4.9 \times 10^{-4}$ | 0.9 |
| GA1 | YAYVHE | $2.6 \times 10^{6}$ | $2.4 \times 10^{-4}$ | 0.1 |
| GS | SGLLAG | $3.7 \times 10^{5}$ | $2.4 \times 10^{-3}$ | 6.6 |
| GW | FGWSNG | $3.2 \times 10^{5}$ | $1.4 \times 10^{-3}$ | 4.2 |
| GY | YSGGNG | $5.2 \times 10^{5}$ | $1.6 \times 10^{-3}$ | 3.2 |

**Table S3. Kinetic parameters of different Protein G-s binding to Fab$^{LRT}$.**

| | $K_{on}$ ($M^{-1}$ $s^{-1}$) | $K_{off}$ ($s^{-1}$) | $K_D$ (nM) | $\chi^2$ ($RU^2$) |
|---|---|---|---|---|
| **G-Fc** | $2.9 \times 10^{4}$ | $1.1 \times 10^{-4}$ | 3.8 | 0.2 |
| **G-Fc2** | $1.1 \times 10^{5}$ | $3.6 \times 10^{-3}$ | 33.7 | 0.5 |
| **G-Fc3** | $1.3 \times 10^{4}$ | $1.4 \times 10^{-4}$ | 10 | 0.7 |

**Table S5. Kinetic parameters of Protein G-Fc binding to Human Fc.**

**Table S5. Sequences**

**PGA1_SNAP_H6 (pEKD40)**
MASTMDIKLTGEFAMDKDCEMKRTTLDSPLGKLELSGCEQGLHEIKLLGKGTSAADAV
EVPAPAAVLGGPEPLMQATAWLNAYFHQPEAIEEFPVPALHHPVFQQESFTRQVLWKLL
KVVKFGEVISYQQLAALAGNPAATAAVKTALSGNPVPILIPCHRVVSSSGAVGGYEGGL
AVKEWLLAHEGHRLGKPGLGLVPRGSPAGGSPTPAVTTYKLVINGRTLSGYTTTTAVDA
ATAEKVFKQYAYVHEVDGEWTYDDATKTFTVTEKPEKLGLEHHHHHH

**PGLM_SNAP_H6 (pEKD40)**
MASTMDIKLTGEFAMDKDCEMKRTTLDSPLGKLELSGCEQGLHEIKLLGKGTSAADAV
EVPAPAAVLGGPEPLMQATAWLNAYFHQPEAIEEFPVPALHHPVFQQESFTRQVLWKLL
KVVKFGEVISYQQLAALAGNPAATAAVKTALSGNPVPILIPCHRVVSSSGAVGGYEGGL
AVKEWLLAHEGHRLGKPGLGLVPRGSPAGGSPTPAVTTYKLVINGRTLSGYTTTTAVDA
ATAEKVFKQLGMMRSVDGEWTYDDATKTFTVTEKPEKLGLEHHHHHH

**PGD_SNAP_H6 (pEKD40)**
MASTMDIKLTGEFAMDKDCEMKRTTLDSPLGKLELSGCEQGLHEIKLLGKGTSAADAV
EVPAPAAVLGGPEPLMQATAWLNAYFHQPEAIEEFPVPALHHPVFQQESFTRQVLWKLL
KVVKFGEVISYQQLAALAGNPAATAAVKTALSGNPVPILIPCHRVVSSSGAVGGYEGGL
AVKEWLLAHEGHRLGKPGLGLVPRGSPAGGSPTPAVTTYKLVINGRTLSGYTTTTAVDA
ATAEKVFKQIDMVSSVDGEWTYDDATKTFTVTEKPEKLGLEHHHHHH

**PF_SNAP_H6 (pEKD40)**
MASTMDIKLTGEFAMDKDCEMKRTTLDSPLGKLELSGCEQGLHEIKLLGKGTSAADAV
EVPAPAAVLGGPEPLMQATAWLNAYFHQPEAIEEFPVPALHHPVFQQESFTRQVLWKLL
KVVKFGEVISYQQLAALAGNPAATAAVKTALSGNPVPILIPCHRVVSSSGAVGGYEGGL
AVKEWLLAHEGHRLGKPGLGLVPRGSPAGGSPTPAVTTYKLVINGRTLSGYTTTTAVDA
ATAEKVFKQYAFGNGVDGEWTYDDATKTFTVTEKPEKLGLEHHHHHH

**PGD_H10 (pHFT2)**
MKHHHHHHHHHHSSDYKDDDDKGENLYFQGSTPAVTTYKLVINGRTLSGYTTTTAVDA
ATAEKVFKQIDMVSSVDGEWTYDDATKTFTVTEKPEKL

**PGLM_H10 (pHFT2)**
MKHHHHHHHHHHSSDYKDDDDKGENLYFQGSTPAVTTYKLVINGRTLSGYTTTTAVDA
ATAEKVFKQLGMMRSVDGEWTYDDATKTFTVTEKPEKL

**PGF_H10 (pHFT2)**
MKHHHHHHHHHHSSDYKDDDDKGENLYFQGSTPAVTTYKLVINGRTLSGYTTTTAVDA
ATAEKVFKQYAFGNGVDGEWTYDDATKTFTVTEKPEKL

**PG-FC (pHFT2)**
MKHHHHHHHHHHSSDYKDDDDKGENLYFQGSTPAVTTYKLVINGKTLKGETTTKAVDA
ETAEKAFKQYANVHEVDGEWTYDDATKTFTVTEKPEKL

**PG-FC2 (pHFT2)**
MKHHHHHHHHHHHSSDYKDDDDKGENLYFQGSTPAVTTYKLVINGKTLKGETTTKAVDA
ETAEKAFKQYAYVHEVDGEWTYDDATKTFTVTEKPEKL

**PG-FC3 (pHFT2)**
MKHHHHHHHHHHHSSDYKDDDDKGENLYFQGSTPAVTTYKLVINGKTLKGETTTKAVDA
ETAEKAFKQYANDNEVDGEWTYDDATKTFTVTEKPEKL

**ASF1_H10 (pHFT2)**
MKHHHHHHHHHHHSSDYKDDDDKGENLYFQGSSSIVSLLGIKVLNNPAKFTDPYEFEITF
ECLESLKHDLEWKLTYVGSSRSLDHDQELDSILVGPVPVGVNKFVFSADPPSAELIPAS
ELVSVTVILLSCSYDGREFVRVGYYVNNEYDEEELRENPPAKVQVDHIVRNILAEKPRV
TRFNIVWDNENEGLE

**Asf1 E11 Fab$^{H}$ (RH2.2)**
SDIQMTQSPSSLSASVGDRVTITCRASQSVSSAVAWYQQKPGKAPKLLIYSASSLYSGV
PSRFSGSRSGTDFTLTISSLQPEDFATYYCQQSSDDPITFGQGTKVEIKRTVAAPSVFIF
PPSDEQLKSGTASVVCLLNNFYPREAKVQWKVDNALQSGNSQESVTEQDSKDSTYSL
SSTLTLSKADYEKHKVYACEVTHQGLSSPVTKSFNRGEC/EISEVQLVESGGGLVQPGG
SLRLSCAASGFNISYSSIHWVRQAPGKGLEWVASISSYYGSTYYADSVKGRFTISADTS
KNTAYLQMNSLRAEDTAVYYCARSRGQASWDYWGQGTLVTVSSASTKGPSVFPLAPS
SKSTSGGTAALGCLVKDYFPEPVTVSWNSGALTSGVHTFPAVLQSSGLYSLSSVVTVP
SSSLGTQTYICNVNHKPSNTKVDKKVEPKSCDKTHT

**ASF1 E5 Fab$^{H}$ (RH2.2)**
SDIQMTQSPSSLSASVGDRVTITCRASQSVSSAVAWYQQKPGKAPKLLIYSASSLYSG
VPSRFSGSRSGTDFTLTISSLQPEDFATYYCQQDGWSLITFGQGTKVEIKRTVAAPSVF
IFPPSDEQLKSGTASVVCLLNNFYPREAKVQWKVDNALQSGNSQESVTEQDSKDSTY
SLSSTLTLSKADYEKHKVYACEVTHQGLSSPVTKSFNRGEC/EISEVQLVESGGGLVQP
GGSLRLSCAASGFNVSYYSIHWVRQAPGKGLEWVASIYPYYGSTSYADSVKGRFTISA
DTSKNTAYLQMNSLRAEDTAVYYCARGYGWALDYWGQGTLVTVSSASTKGPSVFPL
APSSKSTSGGTAALGCLVKDYFPEPVTVSWNSGALTSGVHTFPAVLQSSGLYSLSSVV
TVPSSSLGTQTYICNVNHKPSNTKVDKKVEPKSCDKTHT

**hUCHT1 Fab$^{LRT}$ (RH2.2)**
SDIQMTQSPSSLSASVGDRVTITCSASQDIRNYLNWYQQKPGKAPKRWIYYTSRLHSG
VPSRFSGSGSGTDYTLTISSLQPEDFATYYCQQGNTLPWTFGQGTKVEIKRTVAAPSVF
IFPPSDLRTGTASVVCLLNNFYPREAKVQWKVDNALQSGNSQESVTEQDSKDSTYSLS
STLTLSKADYEKHKVYACEVTHQGLSSPVTKSFNRGEC/EISEVQLVESGGGLVQPGGS
LRLSCAASGFNFTGYTIHWVRQAPGKGLEWMGLINPYKGVSTYNQKFKDKATISTDKS
KNTAYLQMNSLRAEDTAVYYCARSGYYGDSDWYFDYWGQGTLVTVSSASTKGPSVFP
LAPSSKSTSGGTAALGCLVKDYFPEPVTVSWNSGALTSGVHTFPAVLQSSGLYSLSSV
VTVPSSSLGTQTYICNVNHKPSNTKVDKKVEPKSCDKTHT

**11M MBP Fab$^{LRT}$ (RH2.2)**

SDIQMTQSPSSLSASVGDRVTITCRASQSVSSAVAWYQQKPGKAPKLLIYSASSLYSG
VPSRFSGSRSGTDFTLTISSLQPEDFATYYCQQASLTALLTFGQGTKVEIKRTVAAPSV
FIFPPSDLRTGTASVVCLLNNFYPREAKVQWKVDNALQSGNSQESVTEQDSKDSTYSL
SSTLTLSKADYEKHKVYACEVTHQGLSSPVTKSFNRGECEISEVQLVESGGGLVQPGG
SLRLSCAASGFNLSSSSIHWVRQAPGKGLEWVASIYSYYGSTSYADSVKGRFTISADT
SKNTAYLQMNSLRAEDTAVYYCAREYHSYWSYSWWPRVGLDYWGQGTLVTVSSAST
KGPSVFPLAPSSKSTSGGTAALGCLVKDYFPEPVTVSWNSGALTSGVHTFPAVLQSS
GLYSLSSVVTVPSSSLGTQTYICNVNHKPSNTKVDKKVEPKSCDKTHT

**Her2 Fab$^{H}$ (pSFV4)**
SDIQMTQSPSSLSASVGDRVTITCRASQDVNTAVAWYQQKPGKAPKLLIYSASFLYSG
VPSRFSGSRSGTDFTLTISSLQPEDFATYYCQQHYTTPPTFGQGTKVEIKRTVAAPSVF
IFPPSDEQLKSGTASVVCLLNNFYPREAKVQWKVDNALQSGNSQESVTEQDSKDSTY
SLSSTLTLSKADYEKHKVYACEVTHQGLSSPVTKSFNRGEC/MKKNIAFLLASMFVFSIA
TNAYAEISEVQLVESGGGLVQPGGSLRLSCAASGFNIKDTYIHWVRQAPGKGLEWVA
RIYPTNGYTRYADSVKGRFTISADTSKNTAYLQMNSLRAEDTAVYYCSRWGGDGFYA
MDYWGQGTLVTVSSASTKGPSVFPLAPSSKSTSGGTAALGCLVKDYFPEPVTVSWNS
GALTSGVHTFPAVLQSSGLYSLSSVVTVPSSSLGTQTYICNVNHKPSNTKVDKKVEPK
SCDKTHT

**EGFR Fab$^{LRT}$ (pSFV4)**
SDIQMTQSPSSLSASVGDRVTITCQASQDISNYLNWYQQKPGKAPKLLIYDASNLETGV
PSRFSGSGSGTDFTFTISSLQPEDIATYFCQHFDHLPLAFGGGTKVEIKRTVAAPSVFIF
PPSDLRTGTASVVCLLNNFYPREAKVQWKVDNALQSGNSQESVTEQDSKDSTYSLSS
TLTLSKADYEKHKVYACEVTHQGLSSPVTKSFNRGEC/QVQLQESGPGLVKPSETLSL
TCTVSGGSVSSGDYYWTWIRQSPGKGLEWIGHIYYSGNTNYNPSLKSRLTISIDTSKT
QFSLKLSSVTAADTAIYYCVRDRVTGAFDIWGQGTMVTVSSAPTKGPSVFPLAPSSKS
TSGGTAALGCLVKDYFPEPVTVSWNSVALTSGVHTFPAVLQSSGLYSLSSVVTVPSSS
LGTQTYICNVNHKPSNTKVDKKVEPKSCDKTHT

**MJ20 Fab$^{H}$ (pSFV4)**
SDIQMTQSPSSLSASVGDRVTITCRASQSVSSAVAWYQQKPGKAPKLLIYSASSLYSGV
PSRFSGSRSGTDFTLTISSLQPEDFATYYCQQSSSSLITFGQGTKVEIKRTVAAPSVFIFP
PSDEQLKSGTASVVCLLNNFYPREAKVQWKVDNALQSGNSQESVTEQDSKDSTYSLS
STLTLSKADYEKHKVYACEVTHQGLSSPVTKSFNRGEC/EISEVQLVESGGGLVQPGGS
LRLSCAASGFNISYSSIHWVRQAPGKGLEWVASIYSYSGYTSYADSVKGRFTISADTSK
NTAYLQMNSLRAEDTAVYYCARSYWYHVGSWHYTGMDYWGQGTLVTVSSASTKGPS
VFPLAPSSKSTSGGTAALGCLVKDYFPEPVTVSWNSGALTSGVHTFPAVLQSSGLYSLS
SVVTVPSSSLGTQTYICNVNHKPSNTKVDKKVEPKSCDKTHT

**PGA1-hFc (pSCSTa)**
TPAVTTYKLVINGRTLSGYTTTTAVDAATAEKVFKQYAYVHEVDGEWTYDDATKTFTVT
EKPEKLGGGGSGGGSGGGGSGGGGSGGGGSSSGSSCPPCPAPELLGGPSVFLFPP
KPKDTLMISRTPEVTCVVVDVSHEDPEVKFNWYVDGVEVHNAKTKPREEQYNSTYRV
VSVLTVLHQDWLNGKEYKCKVSNKALPAPIEKTISKAKGQPREPQVYTLPPSREEMTK

NQVSLTCLVKGFYPSDIAVEWESNGQPENNYKTTPPVLDSDGSFFLYSKLTVDKSRW
QQGNVFSCSVMHEALHNHYTQKSLSLSPGK

**PGLM-hFc (pSCSTa)**
TPAVTTYKLVINGRTLSGYTTTTAVDAATAEKVFKQLGMMRSVDGEWTYDDATKTFTV
TEKPEKLGGGGSGGGSGGGGSGGGGSGGGGSSSGSSCPPCPAPELLGGPSVFLFP
PKPKDTLMISRTPEVTCVVVDVSHEDPEVKFNWYVDGVEVHNAKTKPREEQYNSTYR
VVSVLTVLHQDWLNGKEYKCKVSNKALPAPIEKTISKAKGQPREPQVYTLPPSREEMT
KNQVSLTCLVKGFYPSDIAVEWESNGQPENNYKTTPPVLDSDGSFFLYSKLTVDKSR
WQQGNVFSCSVMHEALHNHYTQKSLSLSPGK

**PGA1-mFc (pSCSTa)**
TPAVTTYKLVINGRTLSGYTTTTAVDAATAEKVFKQYAYVHEVDGEWTYDDATKTFTVT
EKPEKLGGGGGGSGGGGSGGGGSCPPCKCPAPNLLGGPSVFIFPPKIKDVLMISLSPI
VTCVVVDVSEDDPDVQISWFVNNVEVHTAQTQTHREDYNSTLRVVSALPIQHQDWMS
GKEFKCKVNNKDLPAPIERTISKPKGSVRAPQVYVLPPPEEEMTKKQVTLTCMVTDFM
PEDIYVEWTNNGKTELNYKNTEPVLDSDGSYFMYSKLRVEKKNWVERNSYSCSVVHE
GLHNHHTTKSFSRTPGK

**PGLM-mFc (pSCSTa)**
TPAVTTYKLVINGRTLSGYTTTTAVDAATAEKVFKQLGMMRSVDGEWTYDDATKTFTV
TEKPEKLGGGGGGSGGGGSGGGGSCPPCKCPAPNLLGGPSVFIFPPKIKDVLMISLS
PIVTCVVVDVSEDDPDVQISWFVNNVEVHTAQTQTHREDYNSTLRVVSALPIQHQDW
MSGKEFKCKVNNKDLPAPIERTISKPKGSVRAPQVYVLPPPEEEMTKKQVTLTCMVTD
FMPEDIYVEWTNNGKTELNYKNTEPVLDSDGSYFMYSKLRVEKKNWVERNSYSCSVV
HEGLHNHHTTKSFSRTPGK
